# Bayesian inference for copy number intra-tumoral heterogeneity from single-cell RNA-sequencing data

**DOI:** 10.1101/2023.10.22.563455

**Authors:** PuXue Qiao, Chun Fung Kwok, Guoqi Qian, Davis J McCarthy

## Abstract

High-resolution molecular characterization of intra-tumoral clonal structure defined by genomic and epigenomic alterations is crucial in understanding the natural history of tumors and advancing cancer treatment strategies. Copy number alterations (CNA) are of notable importance as both drivers and markers of clonal structure that can now be assayed at individual cell resolution. However, specific computational methods are needed for accurate inference of clonal profiles and cell states from sparse and noisy single-cell ’omics data. Here, we develop a new Bayesian model to utilize single-cell RNA sequencing (scRNA-seq) data for automatic analysis of intra-tumoral clonal structure with respect to CNAs, without reliance on prior knowledge. The model clusters cells into sub-tumoral clones while simultaneously identifying CNA events in each clone, jointly modelling input from gene expression and germline single-nucleotide polymorphisms. Unlike previous methods, our approach automatically infers the number of clones present in the tumor. In detailed simulation studies our model frequently achieves very high (>90%) cell clustering accuracy and high (>80%) CN state inference accuracy, even in settings of high variance and sparsity. Overall, our method compares strongly against existing software tools. Application to human metastatic melanoma tumor data demonstrates accurate clustering of tumor and non-tumor cells, and reveals clonal CNA profiles that highlight functional gene expression differences between clones from the same tumor. Our method is implemented in a publicly-available, open-source R package, Chloris.

## 1 Introduction

Copy number alteration (CNA) is a major class of genomic drivers of cancer (Vogelstein et al., 2013), which refers to alterations where certain regions of the genome exhibit an abnormal number of copies (Steele et al., 2022). The process of CNA accumulation during tumor progression gives rise to sub-clones with distinct CN profiles within the same tumor specimen (Figure 1A), known as intratumoral heterogeneity (ITH). ITH poses a key challenge to current standards of cancer treatment (Turajlic et al., 2019), therefore understanding the structure of ITH is crucial for improving cancer prognosis and approaches to treatment (Dagogo-Jack and Shaw, 2018; McCarthy et al., 2020).

**Figure 1.**
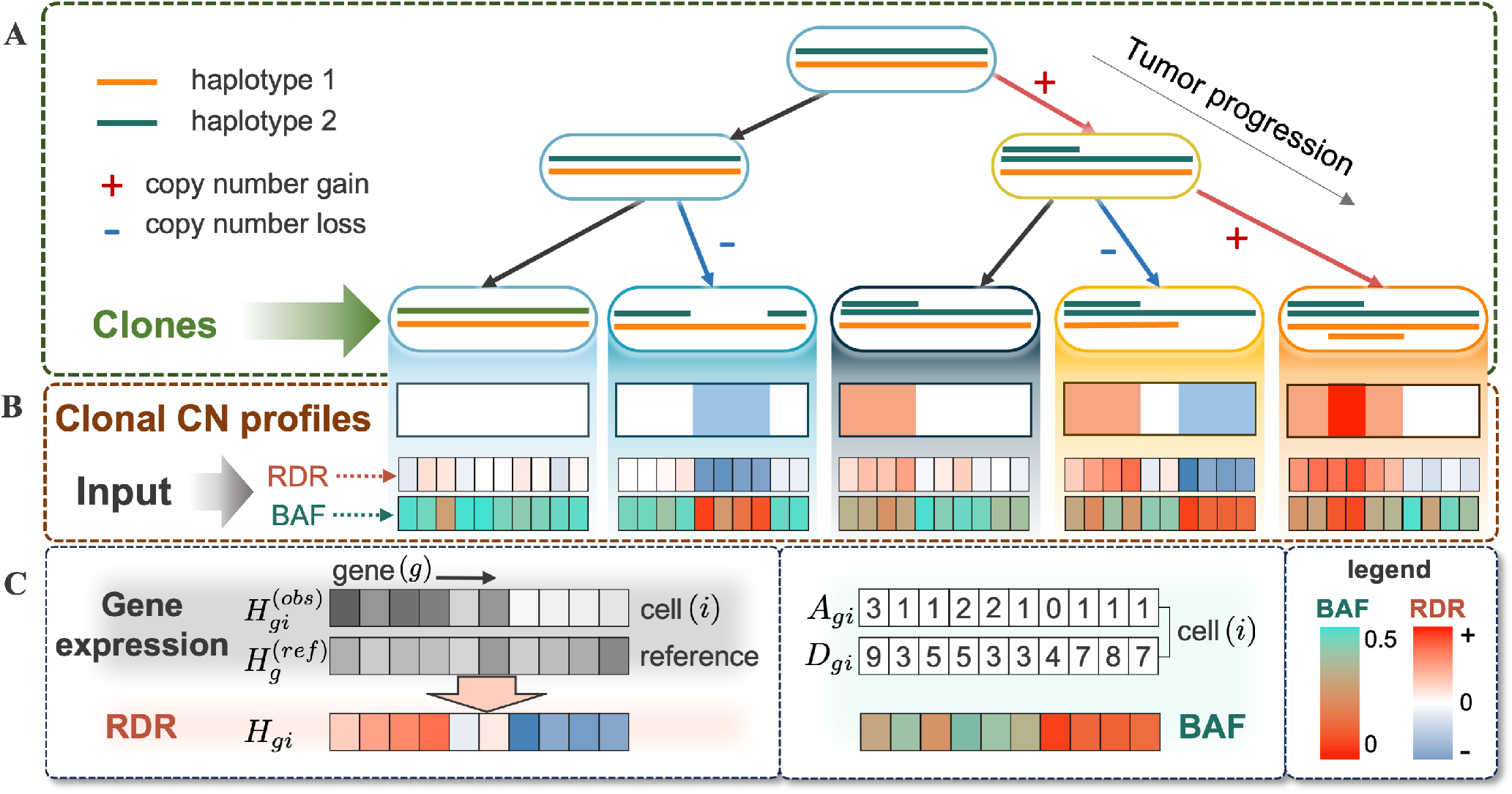
**(A)** An example process of tumor progression: starting from a copy-neutral normal cell, five distinct sub-clones evolve due to the accumulation of copy number gains and losses over time. **(B)** First row: The CN profile representation for each of the clones, where colors indicate four CN states: loss 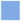, neutral□, gain1 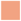 and gain2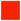 respectively. Second and third row: Stylized example of the associated input (Relative Depth Ratio (RDR) and B-Allelic Fraction (BAF) described in Section 1 and 2.1) from a cell in the according clone, where RDR is loosely correlated with the copy number, while BAF provides an indication of the presence or absence of a CNA event. **(C)** Visual representation of the inputs: The RDR *H*_*gi*_, whichreflects the fluctuation in gene expression of a cell 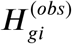 relative to that of reference *H*^(*ref*)^. The BAF(***A***_*gi*_, ***D***_*gi*_), visualized by their ratio ***A***_*gi*_/***D***_*gi*_.

Existing statistical methods for analyzing CNAs include Bayesian models (Lee et al., 2016; Guha et al., 2008), segmentation methods (Olshen et al., 2004; Hu et al., 2016) and penalized regression (Wang and Hu, 2011). However, these methods are designed for traditional bulk sequencing data that averages out genetic information of thousands to millions of cells, masking critical details hidden within cellular heterogeneity.

Single-cell ‘omics technologies have emerged as a powerful tool for assaying individual cells from complex samples. They offer immense potential for studying ITH with unprecedented resolution and pave the way for personalized therapeutic interventions (Saliba et al., 2014; Wen and Tang, 2022). Our focus is to utilize single-cell RNA sequencing (scRNA-seq) data to cluster single cells into clones and infer CN profiles of each clone, which is typically represented by a sequence or vector of CN states (Figure 1B, first row).

Two types of signals related to CNAs are discernible in scRNA-seq data. One involves the gene expression levels. Relative to the levels in a normal diploid cell, the transcriptional expression of a gene affected by CNAs in a tumor cell shifts in proportion to the copy numbers. So the ratio between gene expression in a tumor cell and that in normal reference cells can serve as a surrogate indicator for CN states, denoted the Relative Depth Ratio (RDR; Figure 1C). The drawback of RDR is that it requires gene expression of both the cells of interest (those suspected to have CNAs, that is, tumor cells) and a reference cell group that is free from somatic CNAs, the identification of which typically requires careful sampling of tumor tissue and/or biological expertise for annotating normal cells within a predominantly tumor population. The second source of information is the B-allelic fraction (BAF), which measures the relative abundance of the two alleles observed from the sequenced reads covering a heterozygous single nucleotide polymorphism (SNP). In a copy-neutral region, it is expected that a balanced or similar amount of reads are observed from the two alleles. In other words, the ratio between the counts from one of the alleles (denoted ***A***) and the total counts from both (denoted ***D***) should be around 0.5. Therefore an imbalanced BAF can be seen as an indication of the presence of CNAs (third row in Figure 1B).

Early attempts have been made to identify CNAs from scRNA-seq data. `InferCNV` (Tirosh et al., 2016) implements a pipeline to visualize CNA events in pre-clustered cell groups. CopyKAT (Gao et al., 2021) adopts a Gaussian mixture model to separate CN states reflected in RDR. Fan et al. (2018) are among the first to highlight the usage of BAF in identifying CNAs. CaSpER (Serin Harmanci et al., 2020) proposes to infer the major CN structure from RDR, followed by making local modifications using BAF. Numbat (Gao et al., 2022) models RDR and BAF with an emphasis on haplotype awareness. While these authors successfully demonstrate the ability to infer CNA clonal structure from scRNA-seq data, there are two common issues in their methods: (1) the partition of cell populations and CNA inference are carried out separately, ignoring the fact that reliable identification of sub-clonal CNA events depends on the accurate partitioning of clonal cell populations, and (2) the number of confident sub-clones relies on manual inspection.

We propose a Bayesian model that provides a unified framework to simultaneously partition single cells into clones and identify CN profiles for each clone, allowing the clonal CN profiles and cell clustering to borrow strength from each other through iterative updates. Specifically, we first develop an observation model to associate the input RDR and BAF to both the latent variables that represent clonal CN states per gene and indicator variables that denote cell clustering. We then construct a latent model that relies on a first-order HMM to capture dependencies between CN states of adjacent genes. The proposed approach is highly automated and relies on essentially no prior knowledge. It suggests the optimal number of clones and identifies the normal reference group. The inference of the posterior distribution is performed by Gibbs sampling and implemented as an R package, Chloris.

The remainder of this article is structured as follows. Section 2 presents the proposed Bayesian model and posterior inference. Section 3 presents controlled simulation studies evaluating the performance of our model relative to existing methods, and Section 4 illustrates the model’s real-life application through publicly accessible data from a melanoma patient.

## 2 Model specification

### 2.1 Data

Let *i* = 1, …, *N* be the index of single cells and *g* = 1, …, *G* be that of ordered genes on a given chromosome shared by all cells. For each gene *g*, we observe gene expression levels in each single cell in a tumor sample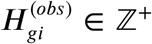 *∈* ℤ^+^ and those in a reference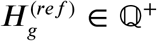. The reference captures gene expression variation in a normal cell unaffected by CNAs. Therefore, the ratio between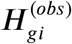 and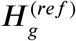 would characterize fluctuations primarily caused by CNAs and can be regarded as a proxy for copy number states. First consider an idealized scenario, in which there is no normalization or measurement error and all cells in a tumor sample exhibit no variation in cell type or cell state. The ratio between the expression level of a gene in a tumor cell and that in a normal cell (or reference) should equal exactly half the number of copies, since a normal cell always has two copies. This ratio on the log_2_ scale, called relative depth ratio (RDR), should then be log_2_ 2/2 = 0 for copy-neutral genes, log_2_ 1/2 = −1 for single-copy losses and log_2_ 3/2 = 0.58 for single-copy gains (second row in Figure 1B).

We then denote the Relative Depth Ratio (RDR) as

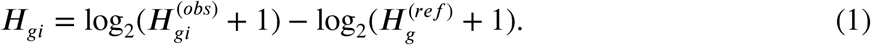

The log transformation of gene count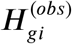 stabilizes the variance in scRNA-seq data, and the pseudo count 1 avoids undefined values while implicitly adding some regularization for small values of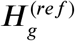.

While the gene expression of suspected tumor cells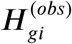 is available from data, that of a reference group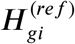 is usually not. An ideal reference would be a matched normal sample from the same patient. Unfortunately, many studies lack appropriate assaying of such cells to provide a reference. Therefore, in Section 2.7, we discuss an alternative way to identify the normal or reference cells within the tumor sample.

The B-allele fraction (BAF) provides another indication of CNAs independent of RDR. It refers to a pair of read counts (***A***_*gi*_, ***D***_*gi*_) for a germline, biallelic, heterozygous SNP *g* in cell *i*, where ***D***_*gi*_ ∈ ℤ^+^ denotes the total count from both alleles, and ***A***_*gi*_ *∈* {0, …, ***D***_*gi*_} represents the counts from the alternative allele.

For SNPs in copy-neutral regions, where there is exactly one copy of each allele, the number of reads from each allele should be similar or balanced. In this case, ***A***_*gi*_ should be around half of ***D***_*gi*_. Whereas in the presence of a CNA, one allele exhibits a higher or lower number of copies relative to the other, leading to an imbalance in allele counts. Without loss of generality, we take *A* as the count of the lower-frequency allele, thus ***A***_*gi*_/***D***_*gi*_ *∈* [0, 0.5] in our analysis. In this article, we do not consider the rare case where both alleles bear copy number alterations in the same gene.

BAF has the advantage of being comparatively free from cell type variations as it does not depend directly on gene expression levels and thus cell types or states. Therefore, inference based on BAF does not rely on the availability of reference cells, but it suffers from much heavier noise than RDR and extreme sparsity, as we can only observe BAF data for SNPs in transcribed regions with sufficient expression, and therefore read coverage, in a given cell.

In our analysis, we use the same index *g* for both genes and SNPs. Even though it is not uncommon to observe no or multiple SNPs within the span of one gene, we aggregate BAF at the gene level to align with RDR for feasibility. Thus, all observations of a certain gene (*g*) in a cell (*i*) can be represented in a vector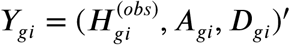, where ^′^ denotes the transpose of a vector.

### 2.2 Cell clustering and clonal CN profiles

The model aims to partition single cells into an unknown number (***K***) of clusters. This partition is represented as a vector of indicators ***I***_*i⋅*_ = (***I***_*i*1_, …, ***I***_*iK*_)^′^, where ***I***_*ik*_ = ***I***(cell *i* belongs to cluster *K*) *∈* {0, 1} and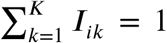 for any *i*. For the rest of this paper, we refer to a cluster of cells as a clone, which could either be a group of normal cells or a tumor sub-clone characterized by CNAs.

Alongside clustering, with the assumption that all cells in one clone share the same CN profile, the model also aims to infer the CN profile for each clone. Denote ***S***_*⋅K*_ = (***S***_1*K*_, …, ***S***_*GK*_) as the CN profile of clone *K*, where *S*_*gK*_ *∈ 𝒮* represents the CN state of gene *g* in clone *K*, and *S* is a pre-specified finite set of CN states. Since RDR and BAF provide independent information on CNA, a CN state is jointly defined by an RDR state, i.e.,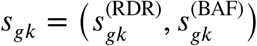, where 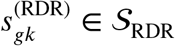 and 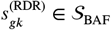 for all *g* and *K*.

We consider four CN states: 𝒮 = {loss, neutral, gain1, gain2}, defined jointly by four RDR states, 𝒮_RDR_ = {−, 0, +, ++}, and two BAF states, 𝒮_BAF_ = {balanced, imbalanced}. Specifically, loss = (-, imbalanced) denotes copy number loss, neutral = (0, balanced) represents copy neutral, and gain1 = (+, imbalanced) and gain2 = (++, imbalanced) refer to single and multiple copy number gain, respectively. These four states are the most commonly observed CNA events in actual tumor samples and are an appropriate level of resolution to be inferred from sparse and noisy single-cell sequence data. The model can be modified to accommodate more states if a higher level of detail is desired for the CN profiles.

Finally, we note that the bijective correspondence between 𝒮_RDR_ and 𝒮 allows us to deduce clonal CNAs to some extent using RDR alone. This feature of the model enables its use in the absence of BAF data. Nonetheless, we show in Section 3.1 that incorporating even sparse BAF data as a complement to RDR data can substantially enhance the model’s performance. For convenience, we refer to the scenario where the input contains only RDR as the “RDR mode”, while we refer to the situation where both RDR and BAF data are available as the “full mode”.

### 2.3 Observation model

We model the RDR (*H*_*gi*_) given state-specific parameters with a Normal distribution, widely adopted for both bulk (Guha et al., 2008) and single-cell data (Tirosh et al., 2016; Fan et al., 2018; Gao et al., 2021). In Section 3.2, we investigate the model’s resilience when this assumption is violated through simulation studies.

We further assume that BAF (***A***_*gi*_ |***D***_*gi*_) follows a Binomial distribution. Thus, given the CN state *s* =(*s*^(RDR)^, *s*^(BAF)^,) we have

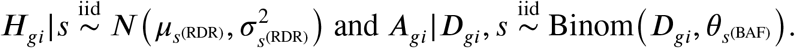

By utilizing the independence between RDR and BAF, and between different cells, the likelihood can be written as

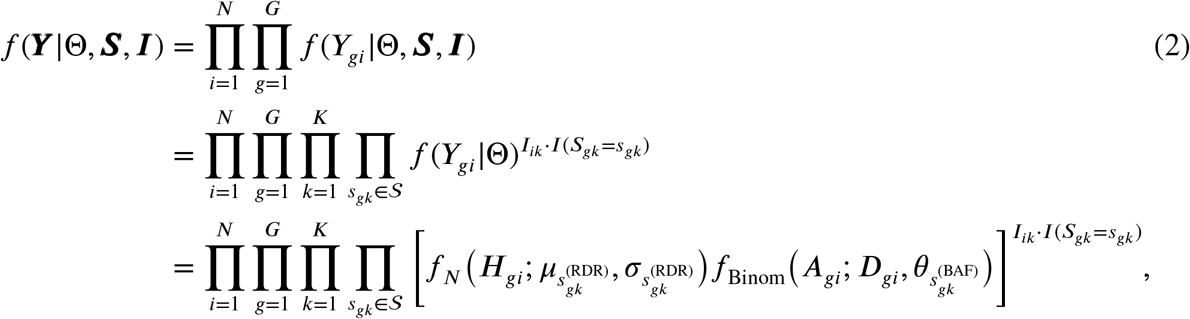

where *f*_*N*_ and *f*_Binom_ de ote the probability density function of Normal and Binomial distribution respectively, and 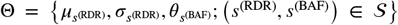 is the set of all state-specific parameters.

### 2.4 Latent model

CNAs are known to carry spatial dependency, meaning that gains or losses at a region/gene are often associated with an increased probability of gains and losses at its neighboring gene. We accommodate this dependency between neighboring genes by building a first-order Hidden Markov Model (HMM; MacDonald and Zucchini, 1997) for each clone *K* independently. Specifically, for any gene index, a Markov model for the CN states assumes that *P* (*S*_*gk*_ = *s*_*gk*_ |*S*_(*g*−1)*k*_, …, *S*_1*K*_)= *P* (*S*_*gk*_ = *s*_*gK*_|*S*_(*g*−1)*k*,_)with conditional probabilities given its neighbouring gene *P* (*S*_*gk*_ = *s*_*gk*_|*S*_(*g*−1)*K*_ = *s*_(*g*−1)*k*)_=*q*_*k*(_*s*_(*g*−1)*k*_, *s*_*gk*)_, where *q*_*k*_(*r, s*), *r, s ∈* 𝒮 denote positive transition probabilities collected in a |𝒮 | × |𝒮| clonal-specific matrix ***Q***_*K*_ . We assume that *S*_0*k*_, the copy number state of the first gene in any clone, follows a flat Categorical distribution. Thus, given ***Q*** = {***Q***_1_, …, ***Q***_*k*_}, the likelihood function for the latent HMM is

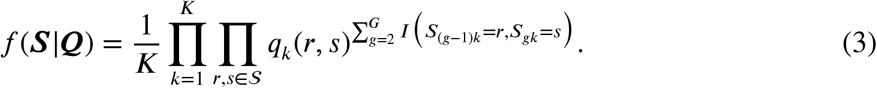

We assume independent Dirichlet priors on ℜ^| 𝒮|^ for the rows of the stochastic matrices ***Q***_*k*_ . That is, with ***Q***_*k*_ (*s, ⋅*) denoting the *s*th row of matrix *Q*_*k*_, we assume that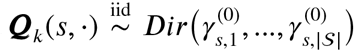 where 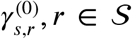 are positive hyperparameters. In our simulation and real data analysis, we fix 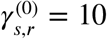 if *s* = *t* and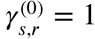 otherwise. In our empirical experience, the choice of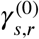 has only a mild effect on the model outcome.

### 2.5 Posterior Inference and label switching handling

Assuming an independent conjugate prior density for emission parameters Θ, the joint prior distribution is

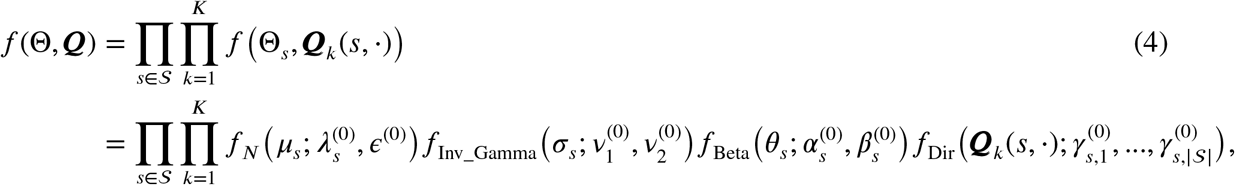

where the informative parameters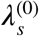 are chosen as the theoretical RDR level for each copy number state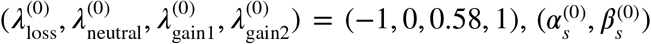 are chosen such that the mean of *θ*_*s*_ equals the theoretical balanced and imbalanced BAF levels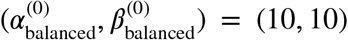 and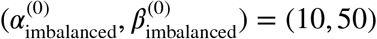 The other parameters are *𝜖*^(0)^ = 0.1,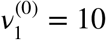, 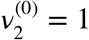.

The joint posterior can thus be written as

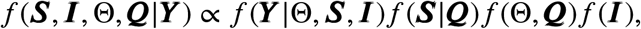

where *f* (*I*) *∼* Categorical(1/*K*, …, 1/*K*) is an uninformative flat prior of cell clustering, priors for other latent variables *f*(Θ, *Q*) are shown in (4), the likelihood function *f* (*Y* Θ, *S, I*) and *f* (*S Q*) are specified in (2) and (3) respectively. Posterior computation given fixed cluster number *K* has been implemented using Gibbs sampling (See Supplementary for details).

A well-established issue in clustering and HMM under the Bayesian framework is the label-switching problem (Scott, 2002). It arises when the likelihood remains invariant under arbitrary permutations of the CN state or clone labels, resulting in inefficient exploration of the posterior by sampling. For the CN state labels, we prohibit any permutation by imposing order constraints on the state-specific parameters, such that *µ*_−_ < *µ*_0_ < *µ*_+_ < *µ*_++_ and *θ*_imbalance_ < *θ*_balance_. To handle potential permutations between clone labels, we incorporate the Equivalence Classes Representatives (ECR) post-process correction method (Papastamoulis and Iliopoulos, 2010) into our MCMC algorithm. We chose ECR because the accuracy of clustering after ECR correction is higher than using other label-switching methods in our experiments (See Supp Figure 2).

### 2.6 Selection of *K*

Although most existing computational tools necessitate a pre-specified value for *K*, it is very difficult to determine the appropriate number of clones without expert examination of the tumor phylogeny and CN profiles. One solution is to select *K* using a suitable model selection criterion, such as the deviance information criterion (DIC; Spiegelhalter et al., 2002). However, applying DIC drastically hinders the efficiency of the algorithm, as it requires repeating the entire MCMC sampling process for each value of *K* within a specified range.

Alternatively, we take advantage of the fact that our model may generate empty clusters when *K* is larger than the actual number of the clusters, and we restart the estimation with the reduced value of *K* when empty clusters appear. Letting the model decide the *K* in this way greatly improves efficiency, as it avoids repeating the MCMC process for the full array of *K*, and also the MCMC process never runs in full for an oversized *K*. Similar ideas have been successfully applied in other clustering algorithms, such as constrained K-means (Bradley et al., 2000) and shrinkage clustering (Hu et al., 2018).

We implement the above by introducing two tuning parameters into the estimation procedure. The pre-specified *𝜔* imposes a constraint on cluster sizes, requiring that all resulting clusters contain at least *𝜔* cells. During the MCMC updates, a cluster will be removed if it contains less than *𝜔* cells for at least *T* consecutive iterations. In all our simulations, we fix *T* = 20 and *𝜔* = 1 (i.e., only empty clusters are removed). Empirically, the clustering result is robust to the specification of *T*, since empty clusters typically appear in the first few iterations and remain empty until the end of the algorithm. In Section 3.1, we evaluate the accuracy of this shrinkage-based method in selecting the right number of clusters in comparison to that of DIC.

### 2.7 Separation of normal and cancer cells

The availability of a high-quality normal reference is crucial to ensure that RDR accurately reflects the underlying copy number states without distortion. An ideal reference would be a bulk or scRNA-seq data sampled from normal tissue of the same patient, with the normal tissue matching the tissue-of-origin (or even more specifically the cell type-of-origin) of the tumor. Ideally, the reference data would be assayed and processed in exactly the same way as the tumor sample under analysis. Unfortunately, such a reference is rarely available.

Therefore, the common practice to date is to separate non-cancer and cancer cells within the available sample. The conventional method is to classify cells according to their cell types using cancer- or immune-specific or other marker genes, then use marker gene expression to define a reference (non-cancer) cell population. In many datasets, the best available option may be to take immune cells as the reference group since immune cells are less prone to carrying somatic mutations than other cells present in a solid tumor. There are several limitations of this method: it requires biological expertise for cell type annotation, immune cells do not always exist in a tumor sample, marker genes can be insufficiently precise, and an immune marker-based approach is inappropriate for any immune cell-derived cancers.

Unlike methods for separating cancer and non-cancer cells based on gene expression, methods based on allelic imbalance of germline SNPs offer appealing advantages. In principle, BAF-based methods for distinguishing cancer from non-cancer cells are less susceptible to sample-to-sample variation and do not require extensive biological expertise, presenting the potential for a fully automated computational approach. Trinh et al. (2022) and Gao et al. (2022) have demonstrated the feasibility of distinguishing normal and cancer cells using BAF signal. Here, we propose a convenient special case of our model that effectively detects the cluster of normal cells. Specifically, a BAF model can be obtained by simply omitting RDR from the full model, with likelihood and prior being

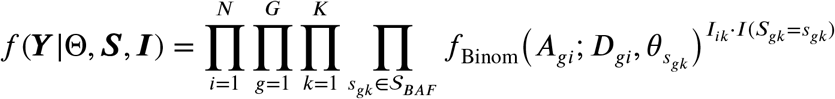

and

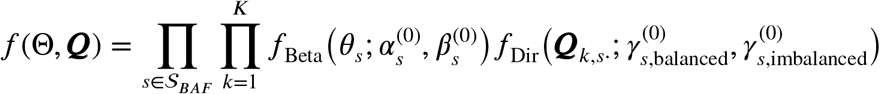

respectively. The cluster *K* where all SNPs are assigned balanced states, that is *X*_*gk*_ = balanced for all *g*, is then identified as the normal cluster.

Despite the inherent sparsity of BAF in single cells, our simulations demonstrate that even with 90% missing data, our model can still identify normal clones with a purity exceeding 90%, meaning that over 90% of the cells in the identified group are genuinely normal (See Supp Figure 1). Finally, note that the BAF model described here is solely used for identifying the reference cell group. It is not advisable to rely on this model for inferring CNA due to the highly sparse nature of BAF data and the fact that BAF does not provide direct evidence regarding the type of CNA, such as gain or loss.

## 3 Simulation studies

We conducted simulations to compare our model’s proficiency with existing methods in executing its three main objectives: (1) clustering cells into clones, (2) identifying CN profiles associated with each clone, and (3) determining the optimal number of clones.

Simulations are executed under two distinct scenarios. The reason for this separation stems from the fact that RDR is not directly observable in real datasets. Instead, it needs to be computed from gene counts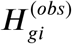 through Equation (1). So the first scenario is a conventional simulation setup, where RDR is generated from a normal distribution based on model assumptions. Whereas in the second scenario, we aim for a closer replicate of the real-world scRNA-seq data by simulating gene counts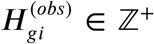. This approach also facilitates comparisons with other software packages, as the majority rely on a gene count matrix as their primary input.

In both scenarios, the CN profile *S*_*⋅K*_ for a tumor clone is generated from independent 4-state Markov chains, with transition probability *q*_*K*_(*s, s*) = 0.98 and ∑_*r*≠*s*_ *q*_*K*_(*s, r*) = 0.02, where *s, r, ∈* 𝒮. Clustering for all cells is generated from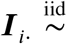 Categorical(1/*K*, …, 1/*K*), where *K* = 4, 5, 6, 7. BAF is generated from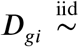 Unif[0, 10] and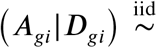 Binom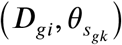 for *k* such that *I*_*ik*_ = 1, where *θ*_balanced_ *∼* Beta(10, 80) and *θ*_imbalanced_ *∼* Beta(10, 15). A setting of 95% missing data (i.e. zero entries in ***A*** and ***D***) is imposed to mimic the sparsity in real data.

### 3.1 Scenario one

In this scenario, RDR given CN state *s*^(RDR)^ is generated from a Normal distribution: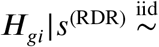 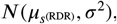, where 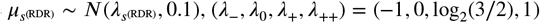 and variance *σ*^2^ = 1, 3, 5 (See examples in Figure 2A).

**Figure 2.**
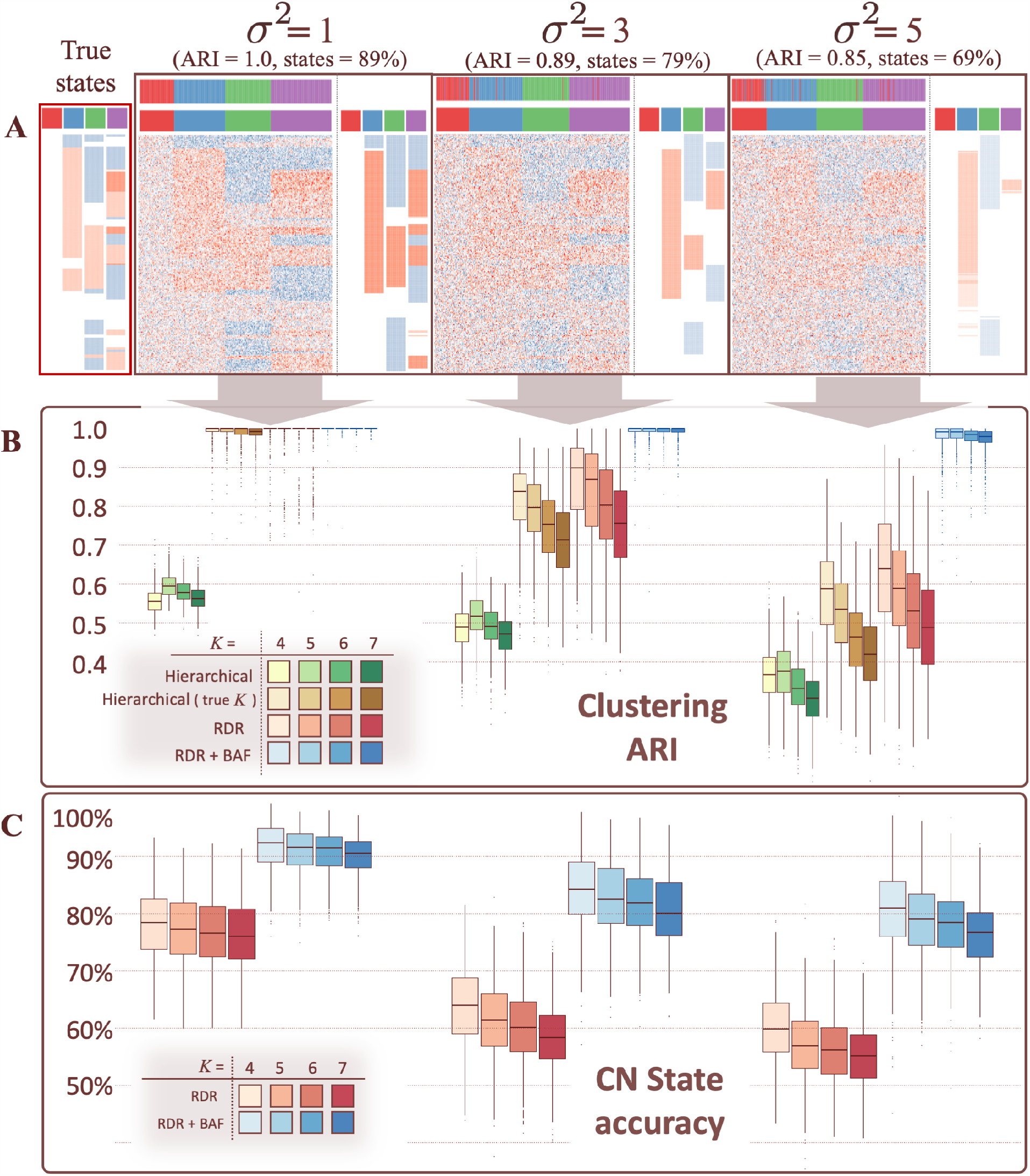
**(A)** Examples of simulated RDR heatmap with variance *σ*^2^ = 1, 3, 5. In the heatmaps, each column and row represents a cell and a gene respectively. Colors 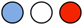 indicate negative, zero and positive values respectively. The color labels on top of the heatmaps show the clustering result using RDR mode (lower), in comparison with the true clustering (upper). The estimated clonal CN profiles Ŝ_*⋅k*_ for the four clones are displayed on the right of each heatmap. **(B)** The Adjusted Rand Index (ARI) comparison between clustering results and simulated truth (y-axis), with increasing variance of RDR (*σ*^2^; x-axis). Red and blue colors represent RDR mode and full mode (RDR + BAF with 95% missing data) respectively, both starting with initial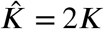. Yellow and green colors represent hierarchical clustering given true *K* and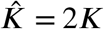. The occupancy of colors indicates the number of clusters *K* = 4, 5, 6, 7. **(C)** Clonal CN state accuracy, measured by the percentage of genes that have been assigned the correct CN state.

In benchmarking cell clustering, we compare our model against the classic hierarchical clustering (Figure 2B), which is used by various computational tools such as `InferCNV` (Tirosh et al., 2016), `HoneyBADGER` (Fan et al., 2018), and Numbat (Gao et al., 2022). One advantage of our model is that it requires only an upper bound of *K* as input instead of the exact *K*. So we provide the initial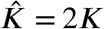 to our model in all simulations. The accuracy of cell clustering is measured using the Adjusted Rand Index (ARI) (Hubert and Arabie, 1985), and it compares the model’s clustering outcome to the simulated ground truth (y-axis). When equipped with the true *K*, hierarchical clustering performs well, only around 0.05 lower in ARI than the RDR mode (red). However, the true *K* is rarely known in practice. When both methods operate with the same starting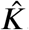, our model strongly outperforms hierarchical clustering (green).

The accuracy for detecting CNA events for each clone is measured by the percentage of genes that have been assigned the correct CN state. Specifically,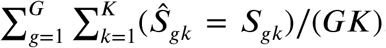 where Ŝ_*gk*_, *S*_*gk*_ ∈ {loss, neutral, gain1, gain2}. For each example in Figure 2A, we show the corresponding estimated clonal CN profiles for the four simulated clones, along with the accuracy of this estimation. In the noisiest simulation (*σ*^2^ = 5), the primary structure within the CN profiles is still largely maintained, only finer details or small CNA regions become obscured or diminished.

We summarize the clustering ARI and state accuracy over 600 simulations (Figure 2B, C). When RDR presents a robust and consistent signal (*σ*^2^ ≤ 1), the “RDR mode” alone is sufficient to produce precise clonal structure, with ARI around 1.0 and state accuracy above 75%. However, in cases where the RDR is noisy, incorporating even extremely sparse BAF with 95% missing data, can significantly boost clustering performance and maintain a 20% increase in state estimation. (see Supp Figure 3 for a more comprehensive result)

Finally, we test the model’s efficacy in determining the optimal number of clones. Here, we compare the performance of three approaches: the shrinkage-based method described in Section 2.6, DIC, and an approximated Bayesian information criterion (BIC; Schwarz, 1978). The two information criteria are approximated as follows:

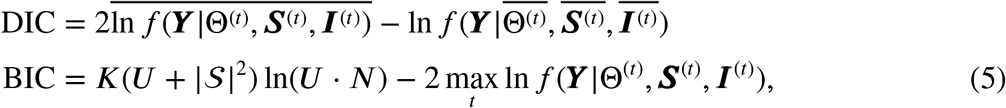

where *f* (*⋅*) is the likelihood function defined in (2), and (Θ^(*t*)^, *S*^(*t*)^, *I*^(*t*)^) are MCMC samples obtained from the *t*th iteration. In obtaining DIC and BIC for a pre-specified *K*, we disable the shrinkage process, meaning that the resulting number of clusters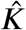 always equals the input value, but there might be empty clusters.

The BIC approximation described in Equation (5) may not equal the true BIC value for two reasons: 1) The second term is not the actual maximum likelihood, but an approximation using the largest likelihood obtained through the MCMC sampling process; 2) In the first term, the number of parameters is estimated by the total number of entries in *X* and *Q* (the only variables whose dimensions vary depending on the value of *K*). However, it is recognized that the exact number of parameters is not clearly defined in a hierarchical Bayesian model. Therefore, the application of BIC here is expected to yield less favorable results than the other methods.

Our simulation results show that BIC manages to select the correct *K* when *σ* is relatively small, but this effectiveness quickly decreases as the RDR becomes noisier. The shrinkage-based method exhibits the greatest resilience to high variance in the RDR, with around 90% of simulation runs successfully recovering the correct *K* when *σ* = 1.1, while less than 70% is reached by DIC under the same condition (see Supp Figure 1).

### 3.2 Scenario two

The results in Scenario one have demonstrated the success of the proposed model given that the input data adheres to the model assumptions. In this section, we aim to evaluate our model using simulation scenarios that much more closely imitate scRNA-seq data from real-world studies.

Specifically, we generate gene counts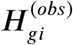 to form a *U* ×*N* expression matrix such that it preserves three key features observed in real datasets: (1) large variation in the library sizes, which represent the difference in sequencing depths between cells, and are typically measured by the total number of counts observed in each single cell, i.e.,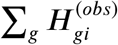 (2) the gene means, or the proportional average gene expression across cells in a biological sample, i.e.,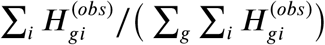 (3) the mean-variance relationship, a widely recognized phenomenon in scRNA-seq data, which refers to the observation that the relationship between the variance of gene expression and the mean expression level is a non-linear function of mean expression. Entries of the expression matrix are generated independently from negative binomial distributions, where parameters are chosen such that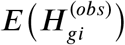 reflects real-world library sizes and gene means (Figure 3A), while Var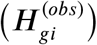 mimics the mean-variance trend of the real data set. The procedure outlined above has been widely accepted in the field as a standard approach for simulating scRNA-seq data. More details can be found in the R package Splatter (Zappia et al., 2017). To incorporate CNA events, we modify the negative binomial parameters such that 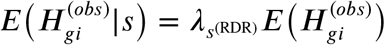, where *λ*_*s*_^(RDR)^ is specified in Section 3.1, and *s* = ∑_*k*_ ***S***_*gk*_***I***_*ik*_ for gene *g* in cell *i*. Given the simulated gene count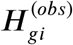 (Figure 3A), the model input RDR *H*_*gi*_ (Figure 3B) can be computed using Equation (1), where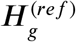 takes the average expression across all normal cells identified from simulated BAF.

**Figure 3.**
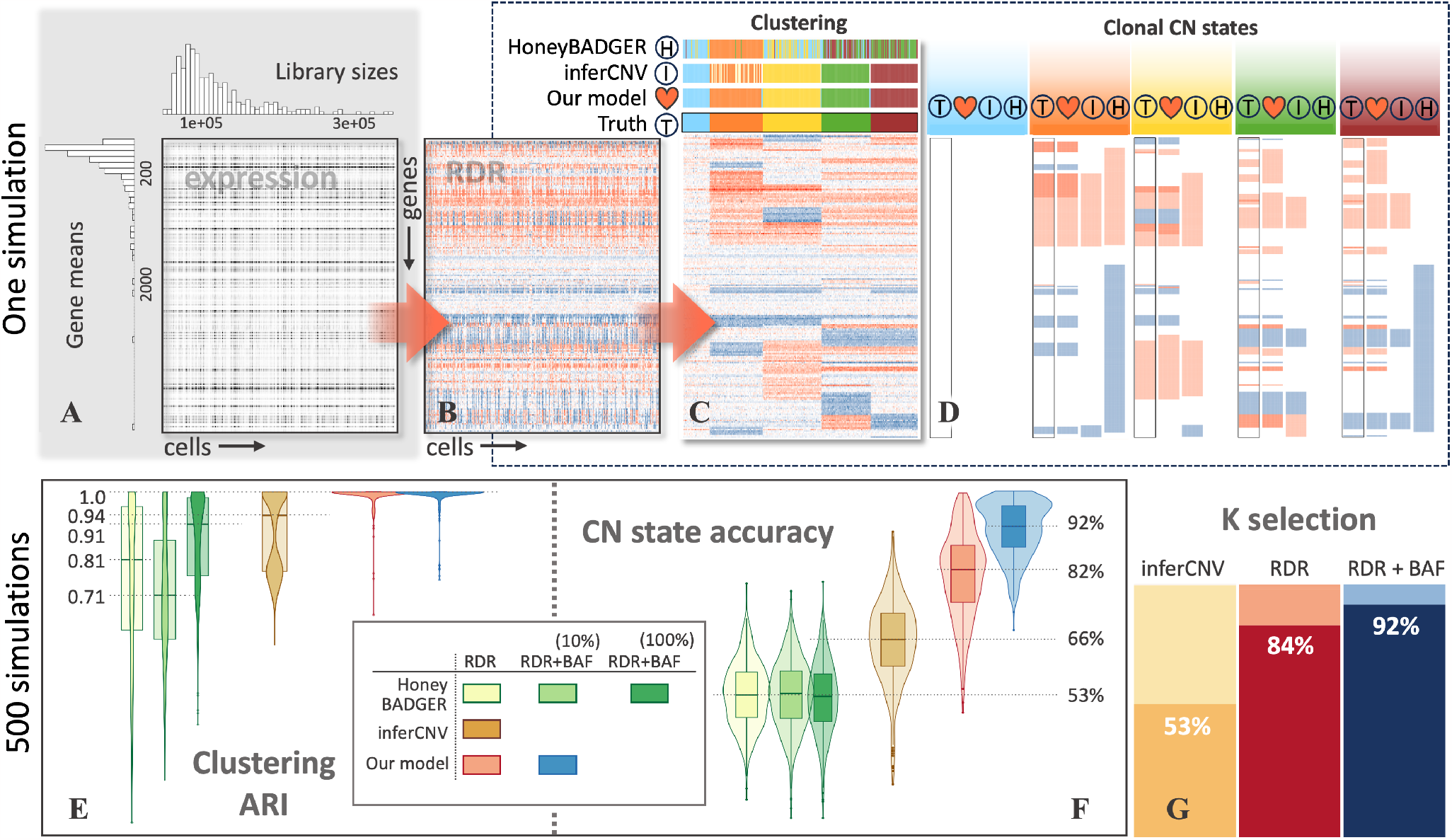
**(A)** An example of simulated gene expression matrix, where entries represent 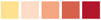 *∈* ℤ^+^, with values indicated by the darkness in the grid. The histograms on the side display the distrib^*g*^u^*i*^ tion of library sizes (column sums) and gene means (row means). Details of the simulation procedure are described in Section 3.2. **(B)** Heatmap of RDR obtained from **(A)**, same format as in Figure 2A. **(C, D)** Results from three methods: our model (heart), `inferCNV` 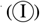 and `HoneyBADGER` 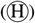, in comparison with the truth 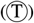. **(C)** Identical RDR as in **(B)**, except that columns/cells are ordered by the true clustering. The clustering results from each method are stacked on top of the heatmap. **(D)** CN profiles for each of the five clones inferred by the three methods. The true profiles are highlighted in black frames. **(E, F)** The clustering ARI and CN state accuracy in 500 simulations summarized in violin and box plots. Three modes of input data: RDR-only for all three models, RDR + 10%BAF (i.e.90% missing data) for our model and `HoneyBADGER`, RDR + 100%BAF (i.e. no missing data) for `HoneyBADGER`. **(G)** The percentage of the 500 simulations in which `inferCNV` and our model (RDR mode and full mode) select the correct number of clusters *K*.

With simulated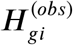 data, we are able to evaluate our model alongside existing computational tools, most of which typically require the gene count matrix as a fundamental input. We consider only software packages that generate comparable results without requiring extra information, therefore excluding Numbat(Gao et al., 2022), which requires phasing information, and `CaSpER` (Serin Harmanci et al., 2020), which does not specify cell partitions. To this end, we focus on two software packages: `inferCNV` (Tirosh et al., 2016), the most widely adopted tool to date but which only considers RDR, and `HoneyBADGER` (Fan et al., 2018), which utilises both RDR and BAF. The primary distinction of our model relative to these tools is that neither `inferCNV` nor `HoneyBADGER` integrates cell clustering and clonal CN profile identification within a unified framework, while ours does. Both `InferCNV` and `HoneyBADGER` first cluster the cells based on RDR using hierarchical clustering. Next, cells belonging to the same cluster are averaged into a single pseudo-bulk vector, resulting in a loss of resolution at cellular level but enabling the convenient application of the classic Viterbi algorithm in the `HiddenMarkov` R package (Harte, 2021). The performance of this two-step procedure heavily depends on the quality of the initial hierarchical clustering output. If cells with distinct CN profiles are erroneously grouped together, they can dilute the signal associated with CNA events.

In every single simulation, the same RDR input is provided to `inferCNV`, `HoneyBADGER`, and our model (Figure 3B). In the presented example (Figure 3C, top), our model correctly recovers both the number of clusters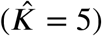 and the cell assignment. `InferCNV` estimates the number of clusters via a non-parametric method similar to that proposed by Kimes et al. (2017), which results in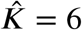 in this example simulation. Although the cluster number is not exact, the clustering remains highly accurate. `HoneyBADGER` requires user input of cluster number, but nevertheless encounters difficulties even with the correct *K* provided. Both our model and `InferCNV` successfully capture the main CN structure for each identified clone, although `InferCNV` achieves a slightly lower resolution (Figure 3D). `HoneyBADGER` is less sensitive to CNA boundaries, resulting in overly large regions for all CN states. One possible explanation is that the state-specific parameters, such as *µ*_*s*_, *σ*_*s*_ and *θ*_*s*_ are treated as fixed constants rather than fitted from the data, thereby losing the flexibility to adapt to the relative fluctuation in RDR specific to each dataset.

We carry out this simulation 500 times and summarize the results in terms of the three model objectives, namely the identification of the true number of clusters, the clustering of cells and the CN profile associated with each cluster. First, we report the percentage of simulations where the models identified the correct *K*. Our model achieves an accuracy of over 80% and `InferCNV` around 50% (Figure 3G). This is also reflected in the ARI clustering metric (Figure 3E), where the ARI of `InferCNV` clustering displays dual peaks: surpassing 0.9 when *K* is correctly identified and falling below 0.8 otherwise.

The full mode operates on BAF with 90% of missing data in conjunction with RDR. Despite the sparsity in BAF, the improvement over RDR-only mode with our model is clear: with both CN state and *K* selection accuracy seeing a 10% increase. In contrast, the full mode of `HoneyBADGER` does not surpass its RDR-only mode. In fact, the clustering with sparse BAF is even worse than using RDR alone (Figure 3E). This is due to HoneyBADGER’s approach of independently handling RDR and BAF, then merging the detected CNA regions from both, in which case sparse or low-quality BAF can yield unfavorable results.

## 4 Real data application

We apply our model to a public scRNA-seq dataset containing 89 single cells from a human metastatic melanoma tumor (Tirosh et al., 2016).

Allele counts of heterozygous germline SNPs (***A*** and ***D***) were generated using cellsnp_lite (Huang and Huang, 2021). SNPs were retained for downstream analyses if they had coverage in at least 5% of the cells (BAF in Figure 4, left). We first identify normal cells using BAF (Section 2.7). The immune cells 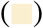, as suggested by immune-specific marker genes, are highly consistent with the identified normal clone (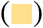), confirming the purity of the reference group (Figure 4, top left).

**Figure 4.**
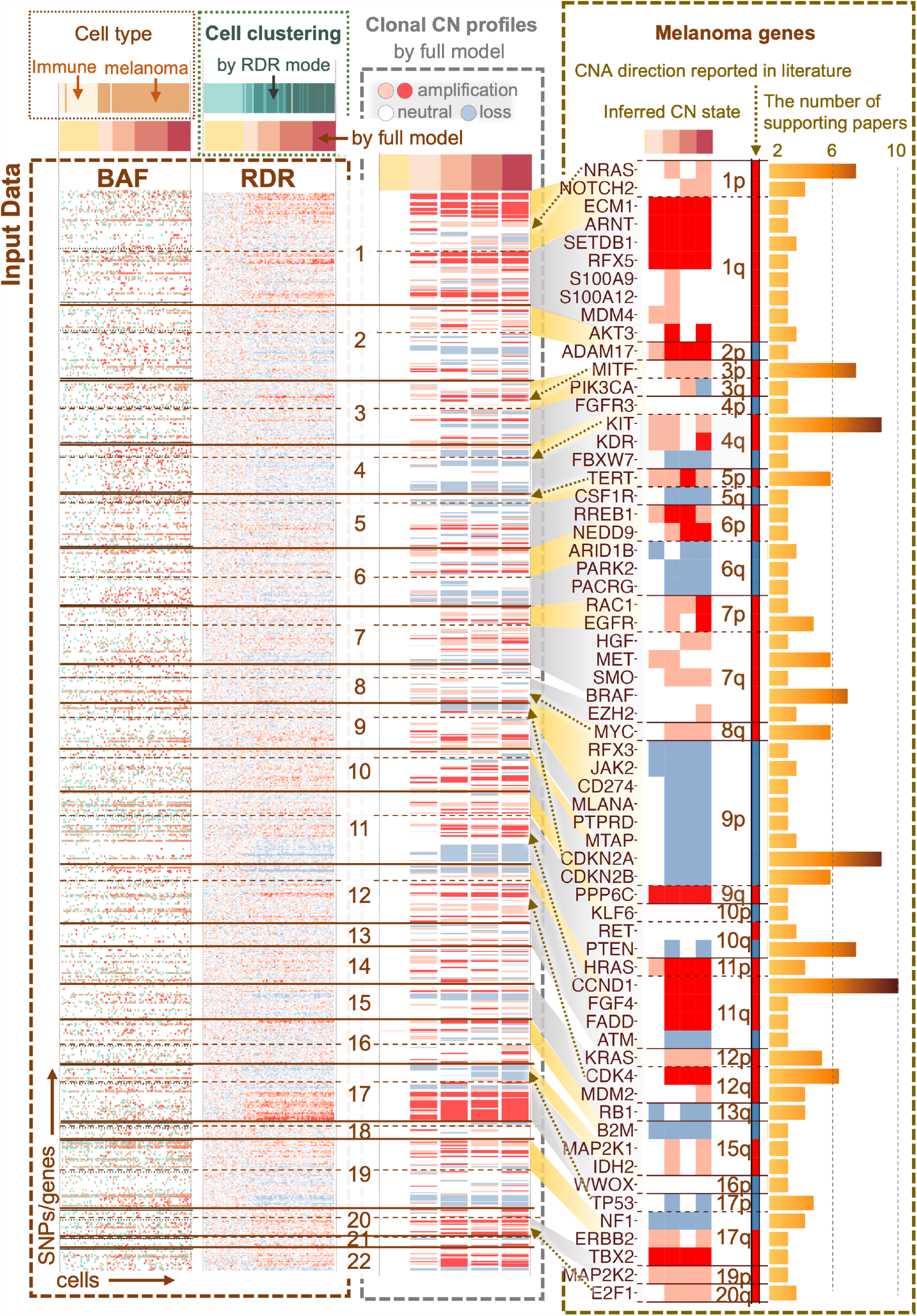
**Left:** Heatmaps of model input, BAF and RDR. Each column and row represents a cell and a gene/SNP respectively. RDR has the same color representation as in Figure 2. BAF entries are colored by the ratio of ***A***_*gi*_/***D***_*gi*_, with turquoise 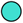 indicating balanced (0.5) and red 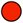 indicating imbalanced (0) ratios, and white representing missing data. The horizontal lines divide chromosomes, and dashed lines marks the positions of the centromere. Columns/cells in both BAF and RDR are ordered by the five cell clusters obtained from the full model (labeled as 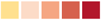). Cell type annotation using marker genes 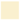 for immune cells and 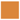 for melanoma cells), and cell clustering using only RDR 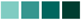 are shown on the top. **Middle:** Inferred clonal CN profiles. **Right:** Consistency of CNAs in 63 known melanoma related genes (gene names listed on the left) between CN states inferred from the model (left) and those reported in 44 melanoma relevant studies (middle, with 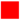 and 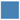 being copy number gain and loss respectively). The number of supporting papers for each gene is shown in bar plot (right).

Standard preprocessing of the gene expression count *H*^(*obs*)^ is done with the scater package (Mc-Carthy et al., 2017). The RDR is then obtained from Equation (1) (Figure 4, left). Genes are ordered according to their genomic coordinates and SNPs are mapped to their overlapping genes. We apply a window smoothing for each cell independently, where the RDR of each gene is smoothed by the weighted sum of nearby genes within a window of size 25 on each side. This approach is a common strategy to overcome sparsity and reduce noise in scRNA-seq data (Tirosh et al., 2016; Fan et al., 2018; Gao et al., 2021).

The full model (with cluster size constraint *𝜔* = 2) converges to five distinct clones 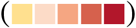 with the first one being the normal clone. The four clusters obtained via the RDR mode 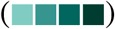 show some resemblance to that from the full model, except for the misclassification of a few melanoma cells into the normal clone. For the rest of this section, we focus on results from the full model.

As expected, the normal clone has been predominantly assigned neutral states in nearly all genes (Figure 4, middle). In observing the CN profiles of the tumor clones, we find many large (chromosomal or arm level) CNA regions consistent with findings reported in existing literature. For example, both Papp et al. (2021) and Wilson et al. (2016) highlight global amplification in 17q and loss in 11q. Pollock et al. (2001) point out that deletions and loss of heterozygosity in 9p has been reported in approximately 60% of melanoma tumors, indicating the possible existence of other melanoma suppressor genes mapping to chromosome 9p other than *CDKN2A*. Large scale chromosome gains in chromosomes 1 and 7, as well as losses in chromosome 4, have also been observed previously (Stark and Hayward, 2007; Moore et al., 2008; Wilson et al., 2016).

We further verify whether the model accurately recovers genes that are widely recognized for harboring CNAs in melanoma. Specifically, we summarize 44 studies on melanoma where CNAs have been observed (Figure 4, right). We record 63 melanoma related genes mentioned in at least two papers. The most commonly reported genes include copy number gains in *NRAS, MITF, KIT, TERT, BRAF, MYC, CCND1, CDK4*, and losses in *CDKN2A, CDKN2B*, and *PTEN*. High consistency has been reached between the inferred CN states in the four sub-tumoral clones and the type of the CNA reported in existing literature.

Finally, we carry out a downstream differential gene expression (DE) analysis to identify genes with any difference in average expression level between each tumor clone and the normal clone (Figure 5), by using the quasi-likelihood *F* -test in the edgeR package (Robinson et al., 2010; Lund et al., 2012) as recommended by Soneson and Robinson (2018). We find hundreds of significantly differentially expressed genes across the four comparisons between tumor and normal clones. From there, we conduct gene set enrichment (pathway) analysis, by applying the camera method (Wu and Smyth, 2012) in the limma package (Smyth, 2004) to the log2-fold-change test statistics obtained from edgeR, to test for enrichment in the 50 Hallmark gene sets from the Molecular Signatures Database (Liberzon et al., 2011). We observe several pathways to be significantly enriched in tumor clones relating to cell proliferation and cancer aggressiveness, like “MYC targets” and “DNA repair” pathways. The inferred clonal CN states of genes in both down- and up-regulated gene sets (Figure 5, windows on each side) are consistent with the enrichment analysis in general.

**Figure 5.**
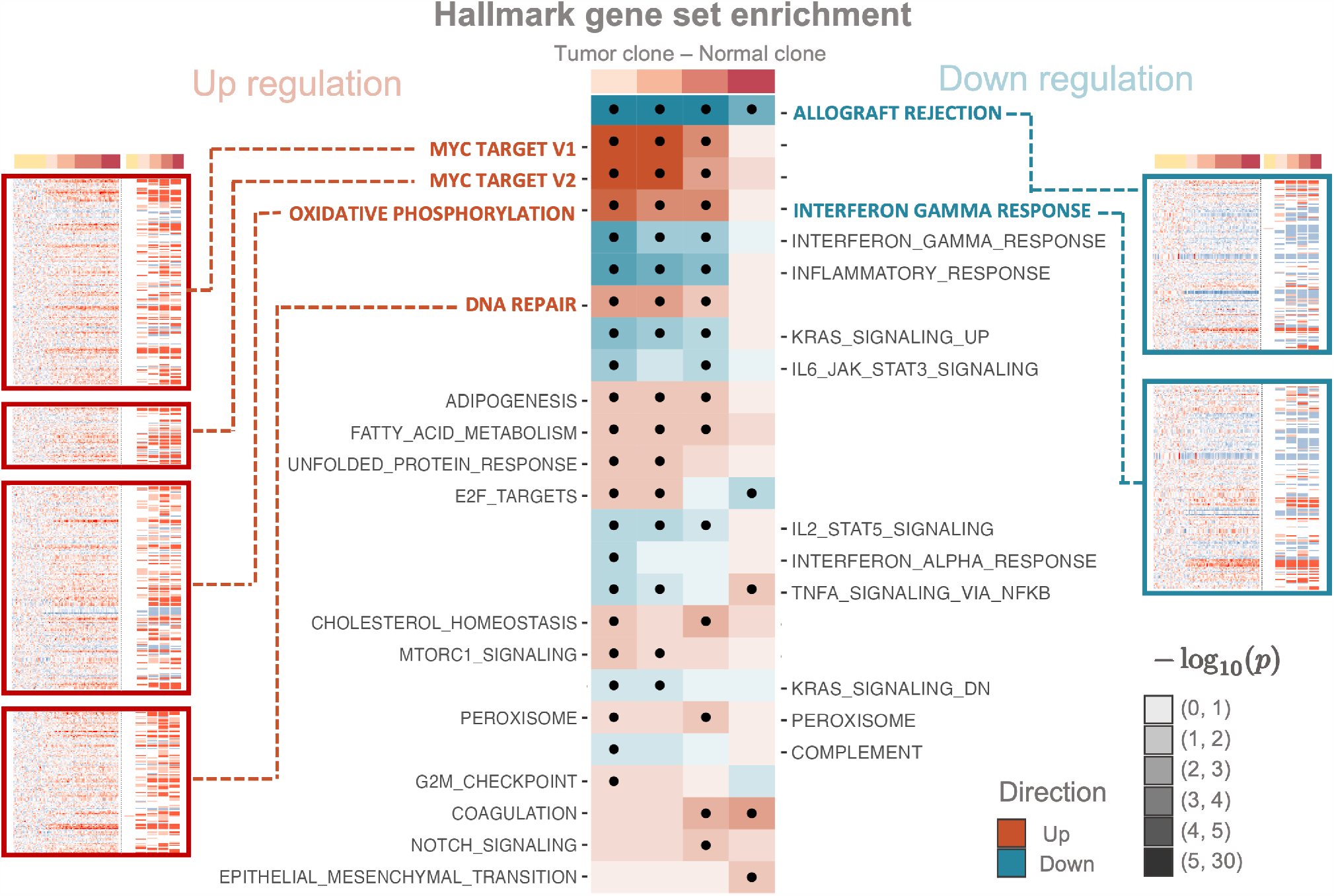
Hallmark gene set enrichment results based on differential gene expression between tumor and normal clones. The heatmap in the center depicts p-values (on log scale) of gene set enrichments (two-sided camera test) for Hallmark gene sets, between the four tumor sub-clones with the normal clone. The colours (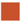 and 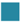) indicate up and down regulation respectively, and dots denote significant enrichment (FDR < 0.05). The windows on the side show RDR alongside inferred clonal CN states of all genes in the top enriched gene sets, with the height of each window proportional to the size of the corresponding gene set.

Collectively, the above results show that our model successfully clusters cells into distinct tumor sub-clones and recovers clonal CNA profiles from scRNA-seq data from human melanoma tissue. Our model’s inferred CN state profiles show excellent agreement with known, common CN changes in 63 genes reported in the literature. Assignment of cells to clones with inferred CN profiles using Chloris thus enables biologically informative downstream analyses of differential expression between clones that can be immediately related back to inferred CN states.

## 5 Discussion

We propose a Bayesian hierarchical approach relying on a hidden Markov model for analyzing intratumoral clonal structure in terms of copy number change. The proposed method infers simultaneously sub-clonal copy number profiles and the partitioning of individual cells into their clone of origin, based on sparse gene expression and germline variant allele data observed in single-cell RNA-sequencing studies.

Reconstructing the single-cell copy number profile from heterogenous cell populations requires the inference of clonal populations and genomic aberrations at the same time. Inference on both aspects in an integrated way is a critical gap in current literature. The ability to do so enables higher sensitivity in discovering candidate genes affected by CNAs that are missed by other algorithms. Our empirical analysis has shown a very favorable comparison with widely adopted software tools `InferCNV` (Tirosh et al., 2016) and `HoneyBADGER` (Fan et al., 2018).

Another strength of our fully automated model is that it relies on essentially no prior knowledge. While many other tools either require an additional input of normal reference cells (Tirosh et al., 2016), or rely on manual inspection to determine the number of sub-clones (Gao et al., 2022), our approach automatically distinguishes normal from cancer cells within the input sample and suggests an optimal number of clusters. In all our analyses, we have used the same default parameterizations specified in Section 2.5. Our simulation studies and empirical experience show that the model outcome is robust to variations in most of the parameters.

Distinguishing normal from cancer cells is a non-trivial prerequisite for accurate CNA inference. We have shown that such classification is feasible by identifying allelic imbalances from single cells, but challenges remain in cases when read coverage of germline SNPs is poor or when normal cells appropriate as a baseline for tumor cells are scarce. Further improvements will be needed to robustly determine the CN baseline in tumors with complex CN profiles.

Another limitation of our model is that it only considers CNA events, and cannot be used to detect other genomic events that contribute to genomic diversity, including chromosomal structural rearrangements, short insertions and deletions (indels) and somatic single-nucleotide variants (SNVs). McCarthy et al. (2020) showed that somatic SNVs can be successfully used for clonal inference and cell-to-clone assignment, and we find it reasonable to assume that indels and complex genomic rearrangements would also be informative identifying clonal structure in tumors. A meaningful extension, therefore, is for our model to incorporate additional sources of genomic mutational information, such as somatic SNVs as a first step, to acquire a more comprehensive mutational profile in tumor sub-clones.

Another valuable extension would be to simultaneously reconstruct the history of copy number events and identify the evolutionary relationships between tumoral clones. Previous studies have succeeded in recovering the phylogenetic clonal tree structure either by incorporating a pre-specified tree as prior (McCarthy et al., 2020), or by facilitating a separate step after the CNA inference (Gao et al., 2022). However, an integrated approach for simultaneously inferring clonal structure and event history reconstruction, like that of Kuipers et al. (2020), for scRNA-seq is yet to be developed.

In summary, we propose a general and automated method that delineates clonal substructure at single cell level and is applicable to high-throughput scRNA-seq data. Its accuracy has been confirmed by both controlled simulations and application to real human melanoma data. Our model’s accurate clonal CN profiles and assignment of cells to clones enable informative downstream analyses of differential gene expression between clones that directly link genomic mutations (CNAs) to the functional behaviour of genomically and transcriptomically heterogeneous cells in tumors. Thus the methods proposed here provide a sharper lens for studying intratumoral heterogeneity in cancer. We hope that the model will serve as a valuable tool for the analysis of genetic and functional states in cancers, and facilitate the development of improved computational approaches in this area.

## Code Availability

The R package Chloris is available at github.com/pqiao29/Chloris. All simulation and analysis are replicable with code and processed data available at github.com/pqiao29/Chloris_experiment.

## Acknowledgments

The authors would like to acknowledge Christina Del Azodi for her extensive help and thoughtful comments on many aspects of this project. St Vincent’s Institute of Medical Research provided IT support and infrastructure used for numerical calculations in this paper enabled via funding from the Victorian State Government Operational Infrastructure Support Program. The authors would like to acknowledge the University of Melbourne for the support and resources provided during the course of this research. This work was supported by funding from the Australian Government through a National Health and Medical Research Council Investigator Grant (GNT1195595) and a National Health and Medical Research Council Project Grant (GNT1162829) awarded to DJM, and by funding from the Baker Foundation awarded to DJM. DJM also acknowledges generous support from Paul Holyoake and Marg Downey.

## References

P. S. Bradley, K. P. Bennett, and A. Demiriz. Constrained K-means clustering. Microsoft Research, Redmond, 20(0):0, 2000.

I. Dagogo-Jack and A. T. Shaw. Tumour heterogeneity and resistance to cancer therapies. Nature Reviews Clinical Oncology, 15(2):81–94, 2018.

J. Fan, H.-O. Lee, S. Lee, D.-e. Ryu, S. Lee, C. Xue, S. J. Kim, K. Kim, N. Barkas, P. J. Park, et al. Linking transcriptional and genetic tumor heterogeneity through allele analysis of single-cell RNA-seq data. Genome Research, 28(8):1217–1227, 2018.

R. Gao, S. Bai, Y. C. Henderson, Y. Lin, A. Schalck, Y. Yan, T. Kumar, M. Hu, E. Sei, A. Davis, et al. Delineating copy number and clonal substructure in human tumors from single-cell transcriptomes. Nature Biotechnology, 39(5):599–608, 2021.

T. Gao, R. Soldatov, H. Sarkar, A. Kurkiewicz, E. Biederstedt, P.-R. Loh, and P. Kharchenko. Haplotype-enhanced inference of somatic copy number profiles from single-cell transcriptomes. bioRxiv, 2022.

S. Guha, Y. Li, and D. Neuberg. Bayesian hidden Markov modeling of array CGH data. Journal of the American Statistical Association, 103(482):485–497, 2008.

D. Harte. HiddenMarkov: Hidden Markov Models. Statistics Research Associates, Wellington, 2021. URL https://www.statsresearch.co.nz/dsh/sslib/. R package version 1. 8-13.

C. W. Hu, H. Li, and A. A. Qutub. Shrinkage Clustering: a fast and size-constrained clustering algorithm for biomedical applications. BMC Bioinformatics, 19(1):1–11, 2018.

J. Hu, L. Zhang, and H. J. Wang. Sequential model selection-based segmentation to detect DNA copy number variation. Biometrics, 72(3):815–826, 2016.

X. Huang and Y. Huang. Cellsnp-lite: an efficient tool for genotyping single cells. Bioinformatics, 37(23):4569–4571, 2021.

L. Hubert and P. Arabie. Comparing partitions. Journal of classification, 2:193–218, 1985.

P. K. Kimes, Y. Liu, D. Neil Hayes, and J. S. Marron. Statistical significance for hierarchical clustering. Biometrics, 73(3):811–821, 2017.

J. Kuipers, M. A. Tuncel, P. Ferreira, K. Jahn, and N. Beerenwinkel. Single-cell copy number calling and event history reconstruction. BioRxiv, pages 2020–04, 2020.

J. Lee, P. Müller, S. Sengupta, K. Gulukota, and Y. Ji. Bayesian inference for intratumour heterogeneity in mutations and copy number variation. Journal of the Royal Statistical Society. Series C, Applied Statistics, 65(4):547, 2016.

A. Liberzon, A. Subramanian, R. Pinchback, H. Thorvaldsdóttir, P. Tamayo, and J. P. Mesirov. Molecular signatures database (MSigDB) 3.0. Bioinformatics, 27(12):1739–1740, 2011.

S. P. Lund, D. Nettleton, D. J. McCarthy, and G. K. Smyth. Detecting differential expression in RNA-sequence data using quasi-likelihood with shrunken dispersion estimates. Statistical applications in genetics and molecular biology, 11(5), Oct. 2012. ISSN 1544-6115. doi: 10.1515/1544-6115.1826. URL 10.1515/1544-6115.1826.

I. L. MacDonald and W. Zucchini. Hidden Markov and other models for discrete-valued time series, volume 110. CRC Press, 1997.

D. J. McCarthy, K. R. Campbell, A. T. Lun, and Q. F. Wills. Scater: pre-processing, quality control, normalization and visualization of single-cell RNA-seq data in R. Bioinformatics, 33(8):1179–1186, 2017.

D. J. McCarthy, R. Rostom, Y. Huang, D. J. Kunz, P. Danecek, M. J. Bonder, T. Hagai, R. Lyu, W. Wang, et al. Cardelino: computational integration of somatic clonal substructure and singlecell transcriptomes. Nature Methods, 17(4):414–421, 2020.

S. R. Moore, D. L. Persons, J. A. Sosman, D. Bobadilla, V. Bedell, D. D. Smith, S. R. Wolman, R. J. Tuthill, J. Moon, V. K. Sondak, et al. Detection of copy number alterations in metastatic melanoma by a DNA fluorescence in situ hybridization probe panel and array comparative genomic hybridization: a southwest oncology group study (s9431). Clinical Cancer Research, 14 (10):2927–2935, 2008.

A. B. Olshen, E. S. Venkatraman, R. Lucito, and M. Wigler. Circular binary segmentation for the analysis of array-based DNA copy number data. Biostatistics, 5(4):557–572, 2004.

P. Papastamoulis and G. Iliopoulos. An artificial allocations based solution to the label switching problem in Bayesian analysis of mixtures of distributions. Journal of Computational and Graphical Statistics, 19(2):313–331, 2010.

O. Papp, V. Doma, J. Gil, G. Markó-Varga, S. Kárpáti, J. Tímár, and L. Vízkeleti. Organ specific copy number variations in visceral metastases of human melanoma. Cancers, 13(23):5984, 2021.

P. M. Pollock, J. Welch, and N. K. Hayward. Evidence for three tumor suppressor loci on chromosome 9p involved in melanoma development. Cancer Research, 61(3):1154–1161, 2001.

M. D. Robinson, D. J. McCarthy, and G. K. Smyth. edgeR: a Bioconductor package for differential expression analysis of digital gene expression data. Bioinformatics, 26(1):139–140, Jan. 2010. ISSN 1367-4803, 1367-4811. doi: 10.1093/bioinformatics/btp616. URL 10.1093/bioinformatics/btp616.

A.-E. Saliba, A. J. Westermann, S. A. Gorski, and J. Vogel. Single-cell RNA-seq: advances and future challenges. Nucleic Acids Research, 42(14):8845–8860, 2014.

G. Schwarz. Estimating the dimension of a model. The Annals of Statistics, pages 461–464, 1978.

S. L. Scott. Bayesian methods for hidden Markov models: Recursive computing in the 21st century. Journal of the American Statistical Association, 97(457):337–351, 2002.

A. Serin Harmanci, A. O. Harmanci, and X. Zhou. CaSpER identifies and visualizes CNV events by integrative analysis of single-cell or bulk RNA-sequencing data. Nature Communications, 11 (1):1–16, 2020.

G. K. Smyth. Linear models and empirical Bayes methods for assessing differential expression in microarray experiments. Statistical Applications in Genetics and Molecular Biology, 3(1), 2004.

C. Soneson and M. D. Robinson. Bias, robustness and scalability in single-cell differential expression analysis. Nature Methods, 15(4):255–261, 2018.

D. J. Spiegelhalter, N. G. Best, B. P. Carlin, and A. Van Der Linde. Bayesian measures of model complexity and fit. Journal of the Royal Statistical Society: Series b (Statistical Methodology), 64(4):583–639, 2002.

M. Stark and N. Hayward. Genome-wide loss of heterozygosity and copy number analysis in melanoma using high-density single-nucleotide polymorphism arrays. Cancer Research, 67(6): 2632–2642, 2007.

C. D. Steele, A. Abbasi, S. A. Islam, A. L. Bowes, A. Khandekar, K. Haase, S. Hames-Fathi, D. Ajayi, A. Verfaillie, P. Dhami, et al. Signatures of copy number alterations in human cancer. Nature, 606(7916):984–991, 2022.

I. Tirosh, B. Izar, S. M. Prakadan, M. H. Wadsworth, D. Treacy, J. J. Trombetta, A. Rotem, C. Rodman, C. Lian, G. Murphy, et al. Dissecting the multicellular ecosystem of metastatic melanoma by single-cell RNA-seq. Science, 352(6282):189–196, 2016.

M. K. Trinh, C. N. Pacyna, G. Kildisiute, C. Thevanesan, A. Piapi, K. Ambridge, N. D. Anderson, E. Khabirova, E. Prigmore, K. Straathof, et al. Precise identification of cancer cells from allelic imbalances in single cell transcriptomes. Communications Biology, 5(1):884, 2022.

S. Turajlic, A. Sottoriva, T. Graham, and C. Swanton. Resolving genetic heterogeneity in cancer. Nature Reviews Genetics, 20(7):404–416, 2019.

B. Vogelstein, N. Papadopoulos, V. E. Velculescu, S. Zhou, L. A. Diaz Jr, and K. W. Kinzler. Cancer genome landscapes. Science, 339(6127):1546–1558, 2013.

H. J. Wang and J. Hu. Identification of Differential Aberrations in Multiple-Sample Array CGH Studies. Biometrics, 67(2):353–362, 2011.

L. Wen and F. Tang. Recent advances in single-cell sequencing technologies. Precision Clinical Medicine, 5(1):pbac002, 2022.

M. A. Wilson, F. Zhao, S. Khare, J. Roszik, S. E. Woodman, K. D’Andrea, B. Wubbenhorst, D. L. Rimm, J. M. Kirkwood, H. M. Kluger, et al. Copy number changes are associated with response to treatment with carboplatin, paclitaxel, and sorafenib in melanoma copy number associated with clinical outcome in melanoma. Clinical Cancer Research, 22(2):374–382, 2016.

D. Wu and G. K. Smyth. Camera: a competitive gene set test accounting for inter-gene correlation. Nucleic Acids Research, 40(17):e133–e133, 2012.

L. Zappia, B. Phipson, and A. Oshlack. Splatter: simulation of single-cell RNA sequencing data. Genome Biology, 18(1):1–15, 2017.

