## Appendix and Supplemental Figures for "Bayesian inference for copy number intra-tumoral heterogeneity from single-cell RNA-sequencing data"

### 1 Gibbs sampler

(1) **Draw $\mu_s$ from**

$$
\begin{aligned}
f(\mu_s|\boldsymbol{Y},\hat{\boldsymbol{S}},\hat{\boldsymbol{I}},\hat{\boldsymbol{Q}},\hat{\Theta}_{-\mu_s},\Theta^{(0)}) &= f(\mu_s|\boldsymbol{H},\hat{\boldsymbol{S}},\hat{\boldsymbol{I}},\hat{\sigma}_s,\lambda_s^{(0)},\epsilon_s^{(0)}) \\
&\propto f(\boldsymbol{H}|\hat{\boldsymbol{S}},\hat{\boldsymbol{I}},\mu_s,\hat{\sigma}_s) f(\mu_s|\lambda_s^{(0)},\epsilon_s^{(0)}),
\end{aligned}
$$

where

$$
\begin{aligned}
f(\boldsymbol{H}|\hat{\boldsymbol{S}},\hat{\boldsymbol{I}},\mu_s,\hat{\sigma}_s) &= \prod_{i=1}^{N}\prod_{k=1}^{K}\prod_{g=1}^{G} f_N(H_{gi};\mu_s,\hat{\sigma}_s)^{\hat{I}_{ik}\cdot I(\hat{S}_{gk}=s)} \\
&\propto \prod_{i=1}^{N}\prod_{k=1}^{K}\prod_{g=1}^{G}\left\{\exp\left[-\frac{1}{2\hat{\sigma}_s}\big(H_{gi}-\mu_s\big)^2\right]\right\}^{\hat{I}_{ik}\cdot I(\hat{S}_{gk}=s)} \\
&= \exp\left[-\frac{1}{2\hat{\sigma}_s}\sum_{i=1}^{N}\sum_{k=1}^{K}\sum_{g=1}^{G}\big(H_{gi}-\mu_s\big)^2\hat{I}_{ik}\cdot I(\hat{S}_{gk}=s)\right] \\
&\propto \exp\Bigg\{-\frac{1}{2\hat{\sigma}_s}\Bigg[\Bigg(\sum_{i=1}^{N}\sum_{k=1}^{K}\sum_{g=1}^{G}\hat{I}_{ik}\cdot I(\hat{S}_{gk}=s)\Bigg)\mu_s^2 \\
&\qquad -2\Bigg(\sum_{i=1}^{N}\sum_{k=1}^{K}\sum_{g=1}^{G} I_{ik}I(\hat{S}_{gk}=s)H_{gi}\Bigg)\mu_s\Bigg]\Bigg\}, \\
f(\mu_s|\lambda_s^{(0)},\epsilon_s^{(0)}) &\propto \exp\left[\frac{1}{2\epsilon_s^{(0)}}(\lambda_s^{(0)}-\mu_s)^2\right].
\end{aligned}
$$

Sample $\hat{\mu}_s$ from

$$
\hat{\mu}_s \sim N(\hat{\lambda}_s,\hat{\epsilon}_s),
$$

where

$$
\begin{aligned}
\hat{\lambda}_s &= \frac{\epsilon_s^{(0)}\Big(\sum_{i=1}^{N}\sum_{k=1}^{K}\sum_{g=1}^{G}\hat{I}_{ik}I(\hat{S}_{gk}=s)H_{gi}\Big)+\hat{\sigma}_s\lambda_s^{(0)}}{\epsilon_s^{(0)}\Big(\sum_{i=1}^{N}\sum_{k=1}^{K}\sum_{g=1}^{G}\hat{I}_{ik}I(\hat{S}_{gk}=s)\Big)+\hat{\sigma}_s}, \\
\hat{\epsilon}_s &= \left(\frac{\sum_{i=1}^{N}\sum_{k=1}^{K}\sum_{g=1}^{G}\hat{I}_{ik}I(\hat{S}_{gk}=s)}{\hat{\sigma}_s}+\frac{1}{\epsilon^{(0)}}\right)^{-1}.
\end{aligned}
$$

(2) **Draw $\sigma_s$ from**

$$
\begin{aligned}
f(\sigma_s|\boldsymbol{Y},\hat{\boldsymbol{S}},\hat{\boldsymbol{I}},\hat{\boldsymbol{Q}},\hat{\Theta}_{-\sigma_s},\Theta^{(0)}) &= f(\sigma_s|\boldsymbol{H},\hat{\boldsymbol{S}},\hat{\boldsymbol{I}},\hat{\mu}_s,\nu_{s,1}^{(0)},\nu_{s,2}^{(0)}) \\
&\propto f(\boldsymbol{H}|\hat{\boldsymbol{S}},\hat{\boldsymbol{I}},\sigma_s,\hat{\mu}_s) f(\sigma_s|\nu_{s,1}^{(0)},\nu_{s,2}^{(0)}),
\end{aligned}
$$

where

$$f(\boldsymbol{H}|\hat{\boldsymbol{S}},\hat{\boldsymbol{I}},\sigma_s,\hat{\mu}_s) = \prod_{i=1}^{N}\prod_{k=1}^{K}\prod_{g=1}^{G} f_N(H_{gi};\hat{\mu}_s,\sigma_s)^{\hat{I}_{ik}\cdot I(\hat{S}_{gk}=s)}$$
$$\propto \prod_{i=1}^{N}\prod_{k=1}^{K}\prod_{g=1}^{G}\left\{\frac{1}{\sigma_s^{1/2}}\exp\left[-\frac{1}{2\sigma_s}\big(H_{gi}-\hat{\mu}_s\big)^2\right]\right\}^{\hat{I}_{ik}\cdot I(\hat{S}_{gk}=s)}$$
$$= \sigma_s^{-\frac{1}{2}\left[\sum_{i=1}^{N}\sum_{k=1}^{K}\sum_{g=1}^{G}\hat{I}_{ik}\cdot I(\hat{S}_{gk}=s)\right]}$$
$$\times\exp\left[-\frac{\sigma_s^{-1}}{2}\sum_{i=1}^{N}\sum_{k=1}^{K}\sum_{g=1}^{G}\hat{I}_{ik}I(\hat{S}_{gk}=s)\big(H_{gi}-\mu_s\big)^2\right],$$
$$f(\sigma_s|\nu_{s,1}^{(0)},\nu_{s,2}^{(0)}) = \sigma_s^{\left(\nu_{s,1}^{(0)}+1\right)}\exp\left(-\sigma_s^{-1}\nu_{s,2}^{(0)}\right).$$

Sample $\hat{\sigma}_s$ from

$$\hat{\sigma}_s \sim \text{Inv-Gamma}(\hat{\nu}_{s,1},\hat{\nu}_{s,2}),$$

where

$$\hat{\nu}_{s,1} = \hat{\nu}_{s,1}^{(0)} + \frac{1}{2}\sum_{i=1}^{N}\sum_{k=1}^{K}\sum_{g=1}^{G} I_{ik}I(\hat{S}_{gk}=s),$$
$$\hat{\nu}_{s,2} = \hat{\nu}_{s,2}^{(0)} + \frac{1}{2}\sum_{i=1}^{N}\sum_{k=1}^{K}\sum_{g=1}^{G} \hat{I}_{ik}I(\hat{S}_{gk}=s)\big(H_{gi}-\mu_s\big)^2$$

(3) **Draw $\theta_s$ from**

$$f(\theta_s|\boldsymbol{Y},\hat{\boldsymbol{S}},\hat{\boldsymbol{I}},\hat{\boldsymbol{Q}},\hat{\Theta}_{-\theta_s},\Theta^{(0)}) = f(\sigma_s|\boldsymbol{A},\boldsymbol{D},\hat{\boldsymbol{S}},\hat{\boldsymbol{I}},\hat{\mu}_s,\alpha_s^{(0)},\beta_s^{(0)})$$
$$\propto f(\boldsymbol{A}|\boldsymbol{D},\hat{\boldsymbol{S}},\hat{\boldsymbol{I}},\theta_s)f(\theta_s|\alpha_s^{(0)},\beta_s^{(0)}),$$

where

$$f(\boldsymbol{A}|\boldsymbol{D},\hat{\boldsymbol{S}},\hat{\boldsymbol{I}},\theta_s) \propto \prod_{i=1}^{N}\prod_{k=1}^{K}\prod_{g=1}^{G}\left[\theta_s^{A_{gi}}(1-\theta_s)^{(D_{gi}-A_{gi})}\right]^{\hat{I}_{ik}I(\hat{S}_{gk}=s)}$$
$$= \theta_s^{\left[\sum_{i=1}^{N}\sum_{k=1}^{K}\sum_{g=1}^{G}A_{gi}\hat{I}_{ik}I(\hat{S}_{gk}=s)\right]}$$
$$\times(1-\theta_s)^{\left[\sum_{i=1}^{N}\sum_{k=1}^{K}\sum_{g=1}^{G}(D_{gi}-A_{gi})\hat{I}_{ik}I(\hat{S}_{gk}=s)\right]},$$
$$f(\theta_s|\alpha_s^{(0)},\beta_s^{(0)}) = \theta_s^{\alpha_s^{(0)}-1}(1-\theta_s)^{\beta_s^{(0)}-1}.$$

Sample $\hat{\theta}_s$ from

$$\hat{\theta}_s \sim \text{Beta}(\hat{\alpha}_s,\hat{\beta}_s),$$

where

$$\hat{\alpha}_s = \alpha_s^{(0)} + \sum_{i=1}^{N}\sum_{k=1}^{K}\sum_{g=1}^{G} A_{gi}\hat{I}_{ik} I(\hat{S}_{gk} = s),$$

$$\hat{\beta}_s = \beta_s^{(0)} + \sum_{i=1}^{N}\sum_{k=1}^{K}\sum_{g=1}^{G} (D_{gi} - A_{gi})\hat{I}_{ik} I(\hat{S}_{gk} = s).$$

(4) **Draw $I_{ik}$ from**

$$P(\hat{\boldsymbol{I}}|\boldsymbol{Y}, \hat{\boldsymbol{X}}, \hat{\boldsymbol{Q}}, \hat{\Theta}, \Theta^{(0)}) \propto f(\boldsymbol{H}|\hat{\boldsymbol{I}}, \hat{\boldsymbol{X}}, \hat{\Theta}) f(\boldsymbol{A}|\boldsymbol{D}, \hat{\boldsymbol{I}}, \hat{\boldsymbol{X}}, \hat{\Theta}) P(\hat{\boldsymbol{I}}|\Theta^{(0)})$$
$$= \prod_{i=1}^{N}\prod_{k=1}^{K}\prod_{s\in\mathcal{S}}\prod_{\substack{g\in\{1,\ldots,G\} \\ s.t. \hat{S}_{gk}=s}} \left[ f_N(H_{gi}; \hat{\mu}_s, \hat{\sigma}_s) f_{\text{binom}}(A_{gi}; D_{gi}, \hat{\theta}_s) \right]^{I_{ik}}$$

Sample $\hat{\boldsymbol{I}_{i.}} = (\hat{I}_{i1}, ..., \hat{I}_{iK})$ from Categorical$(\hat{\varphi}_{i1}, ..., \hat{\varphi}_{iK})$, where

$$\hat{\varphi}_{ik} = \frac{\tilde{\varphi}_{ik}}{\sum_{k=1}^{K} \tilde{\varphi}_{ik}},$$
$$\tilde{\varphi}_{ik} = \exp\left\{ \sum_{s\in\mathcal{S}} \sum_{\substack{g\in\{1,\ldots,G\} \\ s.t. \hat{S}_{gk}=s}} \left[ \log f_N(H_{gi}; \hat{\mu}_s, \hat{\sigma}_s) + \log f_{\text{binom}}(A_{gi}; D_{gi}, \hat{\theta}_s) \right] \right\}.$$

(5) **Sample $S_{gk}$**

We sample $(S_{1k}, ..., S_{Gk})$ for each clone $k$ independently, following the forward-backward Gibbs sampler proposed by Scott (2002):

(5.1) **Forward:**

Let $P_g^{(k)}(r, s) = P(S_{g-1,k} = r, S_{g,k} = s|\boldsymbol{Y}_1^{(k)}, ..., \boldsymbol{Y}_g^{(k)}, \hat{\boldsymbol{Q}}_k, \hat{\Theta})$ for $g = 2, ..., G$, where $\boldsymbol{Y}_g^{(k)} = (Y_{gi})$ for $i$ s.t. $I_{ik} = 1$, which can be estimated as

$$\hat{P}_g^{(k)}(r, s) = \frac{\tilde{P}_g^{(k)}(r, s)}{\sum_{r,s\in\mathcal{S}} \tilde{P}_g^{(k)}(r, s)},$$

where

$$\tilde{P}_g^{(k)}(r, s) = f\big(\boldsymbol{Y}_g^{(k)}|\hat{\Theta}_s\big) \hat{\boldsymbol{Q}}_k(r, s) \hat{\pi}_{g-1}(r),$$
$$f\big(\boldsymbol{Y}_g^{(k)}|\hat{\Theta}_s\big) = \prod_{i=1}^{N} \left[ f_N(H_{gi}; \hat{\mu}_s, \hat{\sigma}_s) f_{\text{binom}}(A_{gi}; D_{gi}, \hat{\theta}_s) \right]^{\hat{I}_{ik}},$$
$$\hat{\pi}_g(s) = \sum_{r\in\mathcal{S}} \hat{P}_g^{(k)}(r, s).$$

(5.2) **Backward:**

Generate $(S_{1k}, ..., S_{Gk})$ from

$$
\begin{aligned}
&P(S_{1k}, ..., S_{Gk}|\boldsymbol{Y}_1^{(k)}, ..., \boldsymbol{Y}_g^{(k)}) \\
&= P(S_{Gk}|\boldsymbol{Y}_1^{(k)}, ..., \boldsymbol{Y}_g^{(k)}) \prod_{g=1}^{G-1} P(S_{(G-g)k}|S_{(G-g+1)k}, ..., S_{Gk}, \boldsymbol{Y}_1^{(k)}, ..., \boldsymbol{Y}_g^{(k)}),
\end{aligned}
$$

where

$$
\begin{aligned}
&P\big(S_{(G-g)k} = r|S_{(G-g+1)k}, ..., S_{Gk}, \boldsymbol{Y}_1^{(k)}, ..., \boldsymbol{Y}_g^{(k)}\big) \\
&= P\big(S_{(G-g)k} = r|S_{(G-g+1)k}, \boldsymbol{Y}_1^{(k)}, ..., \boldsymbol{Y}_g^{(k)}\big) \\
&= \frac{P\big(S_{(G-g)k} = r, S_{(G-g+1)k} = s|\boldsymbol{Y}_1^{(k)}, ..., \boldsymbol{Y}_{G-g+1}^{(k)}, \hat{\boldsymbol{Q}}_k, \hat{\Theta}\big)}{P\big(S_{(G-g+1)k} = s|\boldsymbol{Y}_1^{(k)}, ..., \boldsymbol{Y}_{G-g+1}^{(k)}, \hat{\boldsymbol{Q}}_k, \hat{\Theta}\big)} \\
&= \frac{P_{G-g+1}^{(k)}(r, s)}{\sum_{r \in \mathcal{S}} P_{G-g+1}^{(k)}(r, s)}.
\end{aligned}
$$

(6) **Draw $\boldsymbol{Q}_k$ from**

$$
\begin{aligned}
f(\boldsymbol{Q}_k|\boldsymbol{Y}, \hat{\boldsymbol{S}}, \hat{\boldsymbol{I}}, \hat{\Theta}, \Theta^{(0)}) &= \frac{f(\boldsymbol{Q}_k, \boldsymbol{Y}, \hat{\boldsymbol{S}}, \hat{\boldsymbol{I}}, \hat{\Theta}, \boldsymbol{\gamma}^{(0)})}{f(\boldsymbol{Y}, \hat{\boldsymbol{S}}, \hat{\boldsymbol{I}}, \hat{\Theta}, \boldsymbol{\gamma}^{(0)})} \\
&= \frac{f(\boldsymbol{Y}|\hat{\boldsymbol{S}})P(\hat{\boldsymbol{S}}|\boldsymbol{Q}_k)f(\boldsymbol{Q}_k|\boldsymbol{\gamma}^{(0)})}{f(\boldsymbol{Y}, \hat{\boldsymbol{S}}, \hat{\boldsymbol{I}}, \hat{\Theta}, \boldsymbol{\gamma}^{(0)})} \\
&\propto P(\hat{\boldsymbol{S}}|\boldsymbol{Q}_k)f(\boldsymbol{Q}_k|\boldsymbol{\gamma}^{(0)}) \\
&= \prod_{r \in \mathcal{S}} \prod_{g=1}^{G} P\Big(\hat{S}_{gk}|\hat{S}_{(g-1)k} = r, \boldsymbol{Q}_k(r, \cdot)\Big) f\big(\boldsymbol{Q}_k(r, \cdot)|\boldsymbol{\gamma}^{(0)}\big) \\
&= \prod_{r \in \mathcal{S}} \prod_{g=1}^{G} \left[\prod_{s \in \mathcal{S}} Q_k(r, s)^{I(\hat{S}_{gk}=s, \hat{S}_{(g-1)k}=r)}\right]\left[\prod_{s \in \mathcal{S}} Q_k(r, s)^{\gamma_s^{(0)}-1}\right] \\
&= \prod_{r \in \mathcal{S}} \prod_{s \in \mathcal{S}} Q_k(r, s)^{\left[\gamma_s^{(0)} + \sum_{g=1}^{G} I(\hat{S}_{gk}=s, \hat{S}_{(g-1)k}=r) - 1\right]}
\end{aligned}
$$

Sample $\hat{\boldsymbol{Q}}_k(r, \cdot)$ from $\text{Dir}(\hat{\gamma}_{r1}, ..., \hat{\gamma}_{r|\mathcal{S}|})$, where $\hat{\gamma}_{rs} = \gamma^{(0)} + \sum_{g=1}^{G} I(\hat{S}_{gk} = s, \hat{S}_{(g-1)k} = r)$.

### 2 Supplementary Simulations

#### 2.1 Simulation for normal cell identification

In this section, we evaluate the accuracy of the model special case (Section **??**) in distinguishing normal and cancer cells. We measure this accuracy by the purity of the identified normal clone, that is the percentage of cells that are truly normal cells in the inferred normal cluster. We test this separation under two scenarios: (1) How well does the model tolerate missing data, and (2) how sensitive is the model in recognizing tumor clones that are very similar to a normal clone?

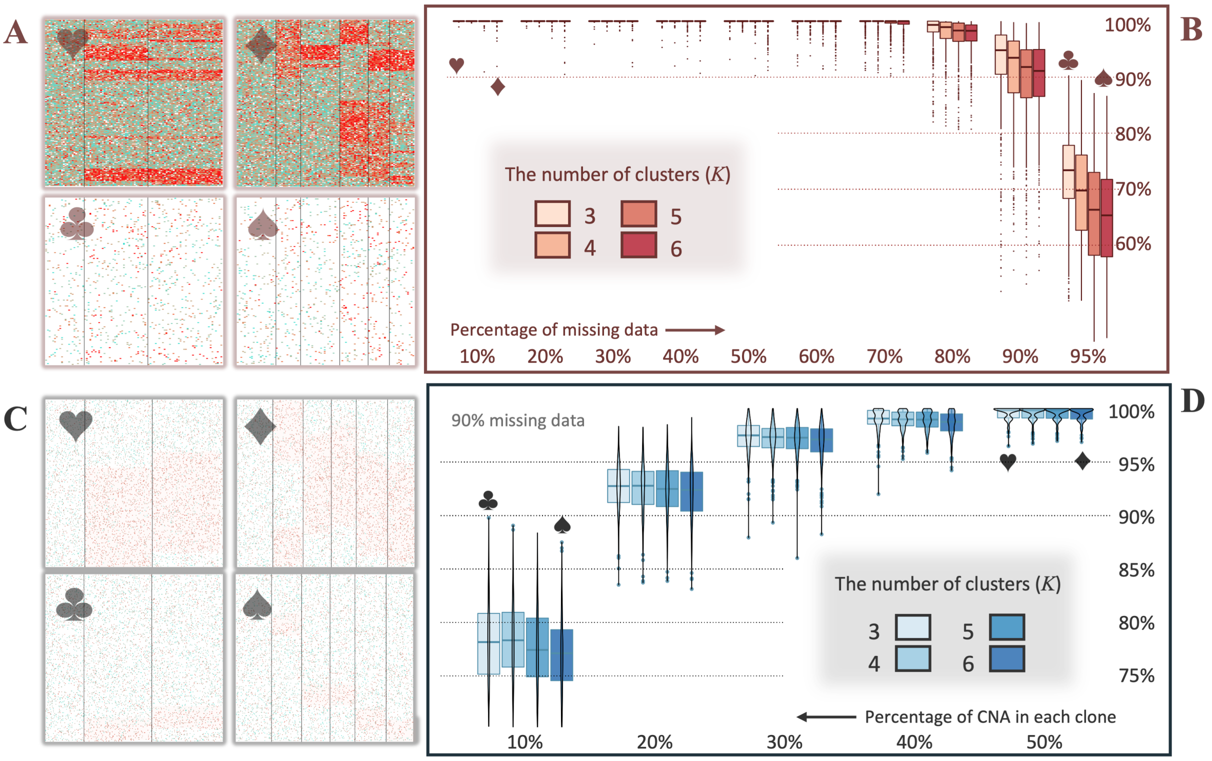

Figure 1: **(A, C)** The heatmap of examples of simulated BAF, with columns and rows indicating cells and SNPs respectively. The black horizontal lines separate the clones. The entries are coloured by the ratio of $A_{gi}/D_{gi}$, with turquoise ● indicating balanced (0.5) and red ● indicating imbalanced (0) ratios, and white representing missing data. **(B, D)** The purity of identified normal cell clone (y-axis), or the percentage of true normal cells in the inferred normal cluster, in 1000 independent simulations. **(B)** The impact of different proportions of missing data (x-axis). **(D)** The impact of different length of CNA regions (x-axis), measured by percentage of SNPs with imbalanced/non-neutral CN state. For convenience, each example is labelled with a card suit symbol. Specifically, in (A) and (B), heart ♥ and diamond ♦ represent BAF with 10% of missing data for 3 and 6 clones respectively, club ♣ and spade ♠ indicate that with 95% of missing data. Similarly, in (C) and (D), heart ♥ and diamond ♦ represent BAF profiles with 100 out of $U = 200$ SNPs affected by CNA in each clone, while club ♣ and spade ♠ indicate those with only 20 imbalanced SNPs.

In both scenarios, we run 1000 independent simulations. In each simulation, we generate $N = 300$ cells and $G = 200$ SNPs. For different levels of clonal complexity, we simulate $K = 3, 4, 5, 6$ clones, among which, clone one is the normal clone (i.e. $S_{g1} =$ balanced for all $g$), while all the rest are tumour sub-clones. Clustering for all cells is generated from $\boldsymbol{I}_{i\cdot} \overset{\text{iid}}{\sim} \text{Categorical}(1/K, ..., 1/K)$. Clonal CN states $S_{gk} \in \mathcal{S}_{\text{BAF}}$ for $k \geq 2$ are simulated differently under the two scenarios, and will be described in the subsequent paragraphs. Total and alternative read counts are generated from $D_{gi} \overset{\text{iid}}{\sim} \text{Unif}[0, 10]$ and $\big(A_{gi}|D_{gi}\big) \overset{\text{iid}}{\sim} \text{Binom}\big(D_{gi}, \theta_{s_{gk}}\big)$, for $k$ such that $I_{ik} = 1$. State specific parameters are generated from $\theta_{\text{balanced}} \sim \text{Beta}(10, 80)$ and $\theta_{\text{imbalanced}} \sim \text{Beta}(10, 15)$.

In scenario one (Figure 1A,B), the CN profile $\boldsymbol{S}_{\cdot k}$ for each tumour clone $k \geq 2$ is generated from independent 2-state Markov chains, with transition probability $q_k(s,s) = 0.98$ and $q_k(s,r) = 0.02$ if $s \neq r$, where $s, r, \in \mathcal{S}_{\text{BAF}}$. We increase the proportion of missing data (i.e. zero entries in $\boldsymbol{A}$ and $\boldsymbol{D}$) from 10% to 95% (symbols ♥♦ and ♣♠ in Figure 1A, respectively). Higher percentages of missing data, as expected, worsen clustering performance (Figure 1B). A more complicated clonal structure (larger $K$) also makes the separation more difficult, especially with sparser data. Nevertheless, the accuracy remains mostly above 90% under around 90% of missing data.

In scenario two, the proportion of missing data is fixed at 90%. We then ask the question: with the presence of 90% missing data, how well does the model perform in identifying tumor clones that closely resemble a normal clone? We quantify this resemblance by the percentage of imbalanced SNPs in a tumour clone, that is $G_{imbal} = \sum_{g=1}^{G} I(X_{gk} = \text{imbalanced})/G$, invariant for all $k \geq 2$. The smaller $G_{imbal}$ a tumour clone has, the more similar it is to a normal clone, and therefore more difficult to differentiate them. So we generate each tumoural CN profile $\boldsymbol{X}_{\cdot k}$ with a controlled length of CNA regions, from $G_{imbal} = 10\%$ (symbols ♣♠ in Figure 1C) to 50% (symbols ♥♦ in Figure 1C). An accuracy of 80% can be reached in the most difficult case where CNAs only affect 20 SNPs; in cases where half of the SNPs exhibit an imbalanced state, this accuracy can reach almost 100% (Figure 1D).

#### 2.2 Label switching handling

Label switching is a well-established issue in clustering and HMM under the Bayesian framework (Scott, 2002). It arises when the likelihood remains invariant under arbitrary permutations of the CN state or clone labels, resulting in inefficient exploration of the posterior by sampling. For the CN state labels, we prohibit any permutation by imposing order constraints on the state-specific parameters, such that $\mu_{-} < \mu_0 < \mu_{+} < \mu_{++}$ and $\theta_{\text{imbalance}} < \theta_{\text{balance}}$. To handle potential permutations between clone labels, we incorporate an existing post-process correction method into our MCMC algorithm. Specifically, we consider three methods: the Equivalence Classes Representatives algorithm (ECR) (Papastamoulis and Iliopoulos, 2010), the geometrically-based Pivotal Reordering Algorithm (PRA) (Marin et al., 2005) and the relabelling algorithm developed by Stephens (2000). All methods can be found in R package `label.switching` (Papastamoulis, 2015).

We compare the performance of the three methods by the clustering accuracy after correction, which is measured by the percentage of cells being correctly assigned to their respective clones (y-axis in Figure 2). For visual convenience, we rank the darkness of the colours according to the performance of the three methods, as well as "none" for not applying any correction, with the darkest and lightest being the best and worst performing method respectively. In our experiments, ECR consistently outperforms the other methods in obtaining accurate clustering results, and is the most memory-efficient option among all. Therefore, we select ECR as the default method to integrate into our

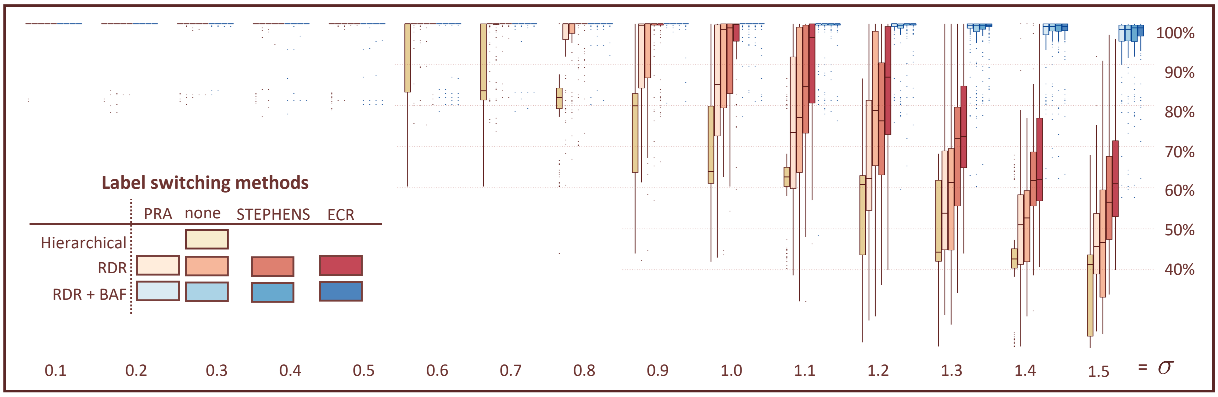

Figure 2: Comparison between label switching correction methods, measured by clustering accuracy after correction (y-axis) with increasing variance in the RDR ($\sigma$; x-axis). Specifically, clustering accuracy is measured by the percentage of cells that have been assigned to the correct cluster. The yellow, red and blue colours indicate hierarchical clustering, RDR mode and full mode respectively. The darkness of the colours indicate the methods considered: ECR, STEPHENS, none (no correction method applied), and PRA. The number of simulated clusters is $K = 5$.

model.

#### 2.3 Full Figure of simulation in Section 3.1

#### 2.4 $K$ selection

We test the model's efficacy in determining the optimal number of clones. Here, we compare the performance of three approaches: the shrinkage-based method described in Section 2.6, DIC and an approximated Bayesian information criterion (BIC; Schwarz, 1978). The two information criteria are approximated as follows:

$$\mathrm{DIC} = 2\overline{\ln f(\boldsymbol{Y}|\Theta^{(t)}, \boldsymbol{S}^{(t)}, \boldsymbol{I}^{(t)})} - \ln f(\boldsymbol{Y}|\overline{\Theta^{(t)}}, \overline{\boldsymbol{S}^{(t)}}, \overline{\boldsymbol{I}^{(t)}})$$

and

$$\mathrm{BIC} = K(U + |\mathcal{S}|^2)\ln(U \cdot N) - 2\max_t \ln f(\boldsymbol{Y}|\Theta^{(t)}, \boldsymbol{S}^{(t)}, \boldsymbol{I}^{(t)}),$$

where $f(\cdot)$ is the likelihood function defined in (2), and $(\Theta^{(t)}, \boldsymbol{S}^{(t)}, \boldsymbol{I}^{(t)})$ are MCMC samples obtained from the $t$th iteration.

We run 500 simulations in both RDR and full mode, with $\sigma$ from 0.1 to 1.5 and true cluster number $K = 5$ (Figure 4). We split and count the simulations according to the estimated number of clusters $\hat{K}$, with the darkest colour indicating the percentage of runs where $\hat{K} = K$ and subsequent colour shades indicating $\hat{K} = K \pm 1, 2, 3$.

BIC manages to select the correct $K$ when $\sigma$ is relatively small, but this effectiveness quickly decreases as the RDR becomes noisier. The shrinkage-based method exhibits the greatest resilience to high variance in the RDR, with around 90% of simulation runs successfully recovering the correct $K$ when $\sigma = 1.1$, while less than 70% is reached by

DIC under the same condition.

### References

J.-M. Marin, K. Mengersen, and C. P. Robert. Bayesian modelling and inference on mixtures of distributions. *Handbook of Statistics*, 25:459–507, 2005.

P. Papastamoulis. label. switching: An R package for dealing with the label switching problem in MCMC outputs. *arXiv preprint arXiv:1503.02271*, 2015.

P. Papastamoulis and G. Iliopoulos. An artificial allocations based solution to the label switching problem in Bayesian analysis of mixtures of distributions. *Journal of Computational and Graphical Statistics*, 19(2):313–331, 2010.

G. Schwarz. Estimating the dimension of a model. *The Annals of Statistics*, pages 461–464, 1978.

S. L. Scott. Bayesian methods for hidden Markov models: Recursive computing in the 21st century. *Journal of the American Statistical Association*, 97(457):337–351, 2002.

M. Stephens. Dealing with label switching in mixture models. *Journal of the Royal Statistical Society: Series B (Statistical Methodology)*, 62(4):795–809, 2000.

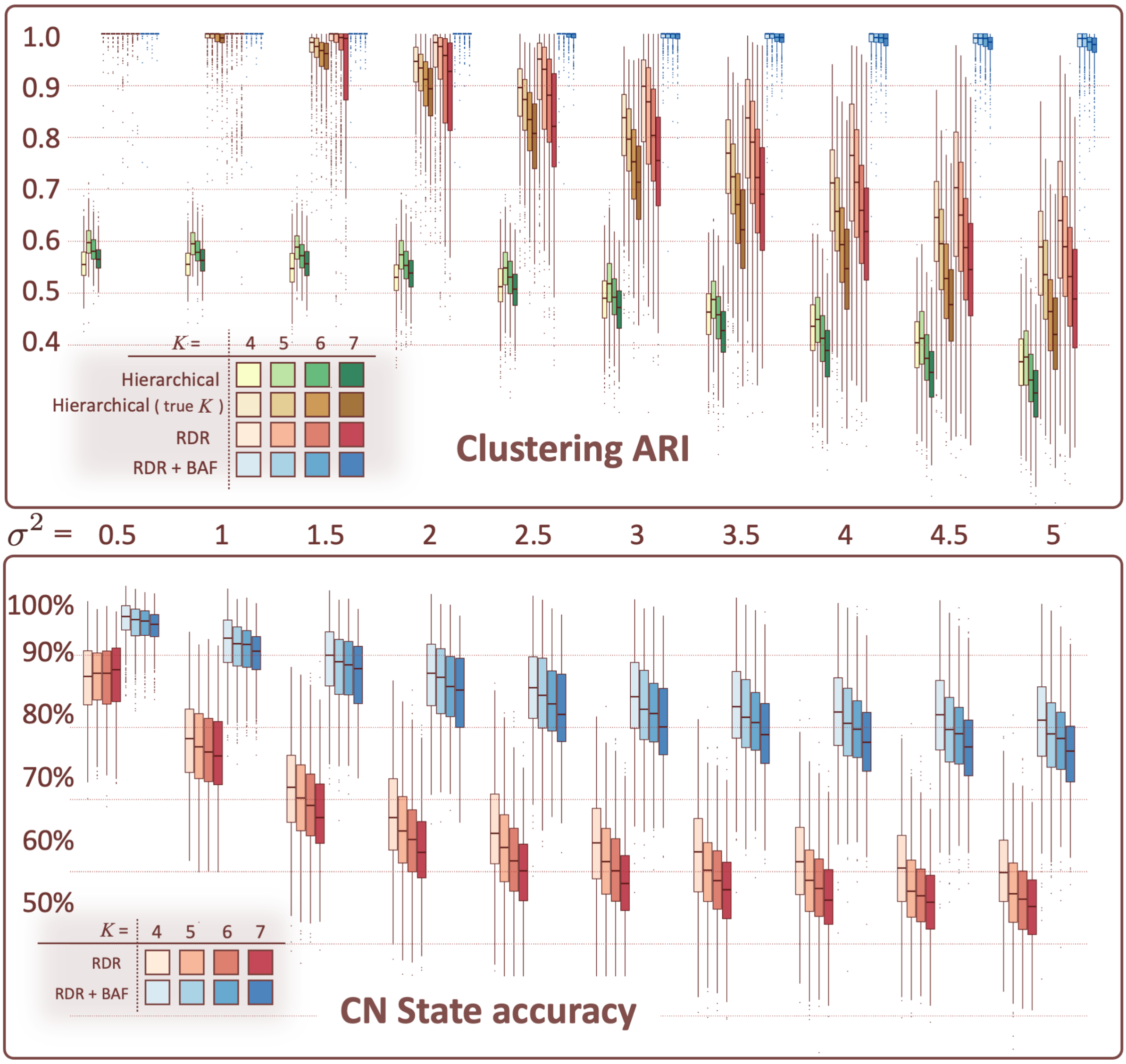

Figure 3: **(A)** The Adjusted Rand Index (ARI) comparison between clustering results and simulated truth (y-axis), with increasing variance of RDR ($\sigma^2 = 0.5, 1, ..., 5$; x-axis). Red and blue colors represent RDR mode and full mode (RDR + BAF with 95% missing data) respectively, both starting with initial $\hat{K} = 2K$. Yellow and green colors represent hierarchical clustering given true $K$ and $\hat{K} = 2K$. The occupancy of colors indicates the number of clusters $K = 4, 5, 6, 7$. **(B)** Clonal CN state accuracy, measured by the percentage of genes that have been assigned the correct CN state.

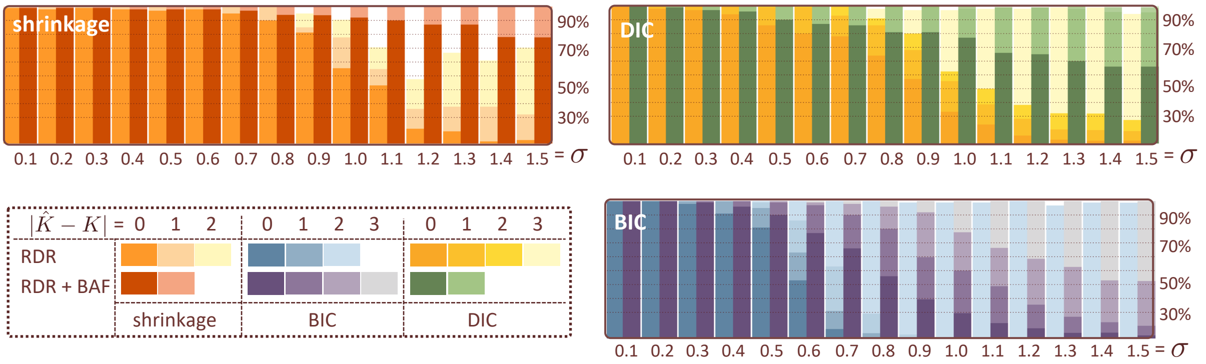

Figure 4: The selection of the optimal number of clusters by three methods: a shrinkage-based method proposed in Section 2.6, deviance information criterion (DIC) and Bayesian information criterion (BIC). The occupancy of the colours in a bar indicates how close the selected $\hat{K}$ is to the true $K$ (i.e. $|\hat{K} - K|$).
